# Neural representations of covert attention across saccades: comparing pattern similarity to shifting and holding attention during fixation

**DOI:** 10.1101/2020.05.10.087270

**Authors:** Xiaoli Zhang, Julie D Golomb

## Abstract

We can focus visuospatial attention by covertly attending to relevant locations, moving our eyes, or both simultaneously. How does shifting versus holding covert attention during fixation compare with maintaining covert attention across saccades? We acquired fMRI data during a combined saccade and covert attention task. On Eyes-fixed trials, participants either held attention at the same initial location (“hold attention”) or shifted attention to another location midway through the trial (“shift attention”). On Eyes-move trials, participants made a saccade midway through the trial, while maintaining attention in one of two reference frames: The “retinotopic attention” condition involved holding attention at a fixation-relative location but shifting to a different screen-centered location, whereas the “spatiotopic attention” condition involved holding attention on the same screen-centered location but shifting relative to fixation. We localized the brain network sensitive to attention shifts (shift > hold attention), and used multivoxel pattern time course analyses to investigate the patterns of brain activity for spatiotopic and retinotopic attention. In the attention shift network, we found transient information about both whether covert shifts were made and whether saccades were executed. Moreover, in the attention shift network, both retinotopic and spatiotopic conditions were represented more similarly to shifting than to holding covert attention. An exploratory searchlight analysis revealed additional regions where spatiotopic was relatively more similar to shifting and retinotopic more to holding. Thus, maintaining retinotopic and spatiotopic attention across saccades may involve different types of updating that vary in similarity to covert attention “hold” and “shift” signals across different regions.

**Significance Statement:** To our knowledge, this study is the first attempt to directly compare human brain activity patterns of covert attention (to a peripheral spatial location) across saccades and during fixation. We applied fMRI multivoxel pattern time course analyses to capture the dynamic changes of activity patterns, with specific focus on the critical timepoints related to attention shifts and saccades. Our findings indicate that both retinotopic and spatiotopic attention across saccades produce patterns of activation similar to “shifting” attention in the brain, even though both tasks could be interpreted as “holding” attention by the participant. The results offer a novel perspective to understand how the brain processes and updates spatial information under different circumstances to fit the needs of various cognitive tasks.

## Introduction

We live in a world with an abundance of visual information, but we have limited visual acuity and cognitive resources. To process visual information across various locations with high sensitivity as needed by daily tasks, we can perform functions like shifting attention allocation covertly or making eye movements. In daily life, covert attention shifts and saccades are often directed to the same to-be-attended location. But we can also covertly attend one location while saccading elsewhere, and the neural mechanisms underlying this case are considerably less explored.

When the eyes are at a stable fixation, covert shifts of attention are often associated with activation in the frontoparietal network (Chica et al., 2013). Specifically, medial superior parietal lobule is activated when covert attention is shifted spatially (Gmeindl et al., 2016; Yantis et al., 2002), between space and feature dimensions (Greenberg et al., 2010), between visual and auditory modalities (Shomstein & Yantis, 2004), and between spatial and nonspatial modalities (Shomstein & Yantis, 2006), suggesting the presence of a general mechanism that mediates shifts of attention.

A number of studies comparing covert attention shifts with overt attention shifts (saccades) further show that these two functions share overlapping brain areas, including intraparietal sulcus (IPS), superior parietal lobule (SPL), and frontal regions like pre-central sulcus/gyrus (Beauchamp et al., 2001; Corbetta et al., 1998; de Haan et al., 2008; Perry & Zeki, 2000). In these neuroimaging studies, a common paradigm is for participants to either shift attention (covert shifts) or make a saccade (overt shifts) between the current fixation point and a target location, with the brain activation in these conditions each contrasted with a baseline condition where no shift happened.

These neuroimaging studies, together with behavioral evidence, suggest a tight coupling between covert spatial attention and eye movements. Covert attentional orientation is an important step preceding saccade execution (Kowler et al., 1995; Peterson et al., 2004). The premotor theory of attention even claims that covert attention simply reflects the central programming of eye movements, just without actual saccade execution (Rizzolatti, Riggio. Dascola, & Umiltá, 1987). However, this theory remains controversial, especially regarding independence between endogenous attention and motor preparation (Smith & Schenk, 2012), and covert spatial attention and saccade target locations can be dissociated in several paradigms, such as anti-saccade tasks (Juan et al., 2004; Smith & Schenk, 2007) and attention in different reference frames, as below.

When attention is allocated to a separate location from the saccade target, the eye movement introduces a discrepancy between retinotopic (eye-centered) and non-retinotopic (e.g. spatiotopic / world-centered) reference frames. Although the spatiotopic reference frame feels more relevant for most behaviors, visual processing starts on our retina in retinotopic coordinates. Behavioral and neural evidence shows that we can allocate attention in both retinotopic and spatiotopic reference frames, though it is debated which is more dominant and whether they differ by brain region (Crespi et al., 2011; Fabius et al., 2016; Fairhall et al., 2017; Golomb et al., 2008; Golomb & Kanwisher, 2012a, 2012b; Melcher & Morrone, 2003; Satel et al., 2012; Shafer-Skelton & Golomb, 2017; Turi & Burr, 2012; Zimmermann et al., 2013).

This ambiguity raises important questions about how our brain processes covert attention across saccades. For example, maintaining covert attention at a stable peripheral real-world location across a saccade would be akin to holding attention in spatiotopic coordinates, but shifting attention in retinotopic coordinates. Here, we take a novel approach to understanding the relationship between covert attention and saccades by comparing the neural patterns associated with holding and shifting covert attention during fixation to retinotopic and spatiotopic attention across saccades. We specifically focus on the attention shift network described above. We acquired fMRI data during a combined saccade and covert attention task, with four critical conditions. On Eyes-fixed trials, participants either held attention at the same initial peripheral location (“hold attention”) or shifted attention to a different location midway through the trial (“shift attention”). On Eyes-move trials, participants made a saccade midway through the trial half of the time, while covertly maintaining either “spatiotopic attention” (hold relative-to-screen, shift relative-to-eyes) or “retinotopic attention” (hold relative-to-eyes, shift relative-to-screen). We used multivoxel pattern time course (MVPTC) analyses to compare whether patterns of brain activity for spatiotopic and retinotopic conditions were more similar to shifting or to holding attention, both in our a priori ROIs, as well as through an exploratory whole-brain searchlight analysis.

## Methods

### Participants

12 right-handed subjects participated in the study (7 females, 5 males, mean age 19.08, range 18-25). An additional left-handed subject was also scanned inadvertently, but the data were not included in our analyses. All subjects reported normal or corrected-to-normal vision. They were pre-screened for MRI eligibility, and they gave informed consent. The study protocol was approved by [anonymous] Institutional Review Board.

### Stimuli and task

The paradigm is shown in Figure 1. Eyes-fixed and Eyes-move tasks were done in separate runs.

**Figure 1.**
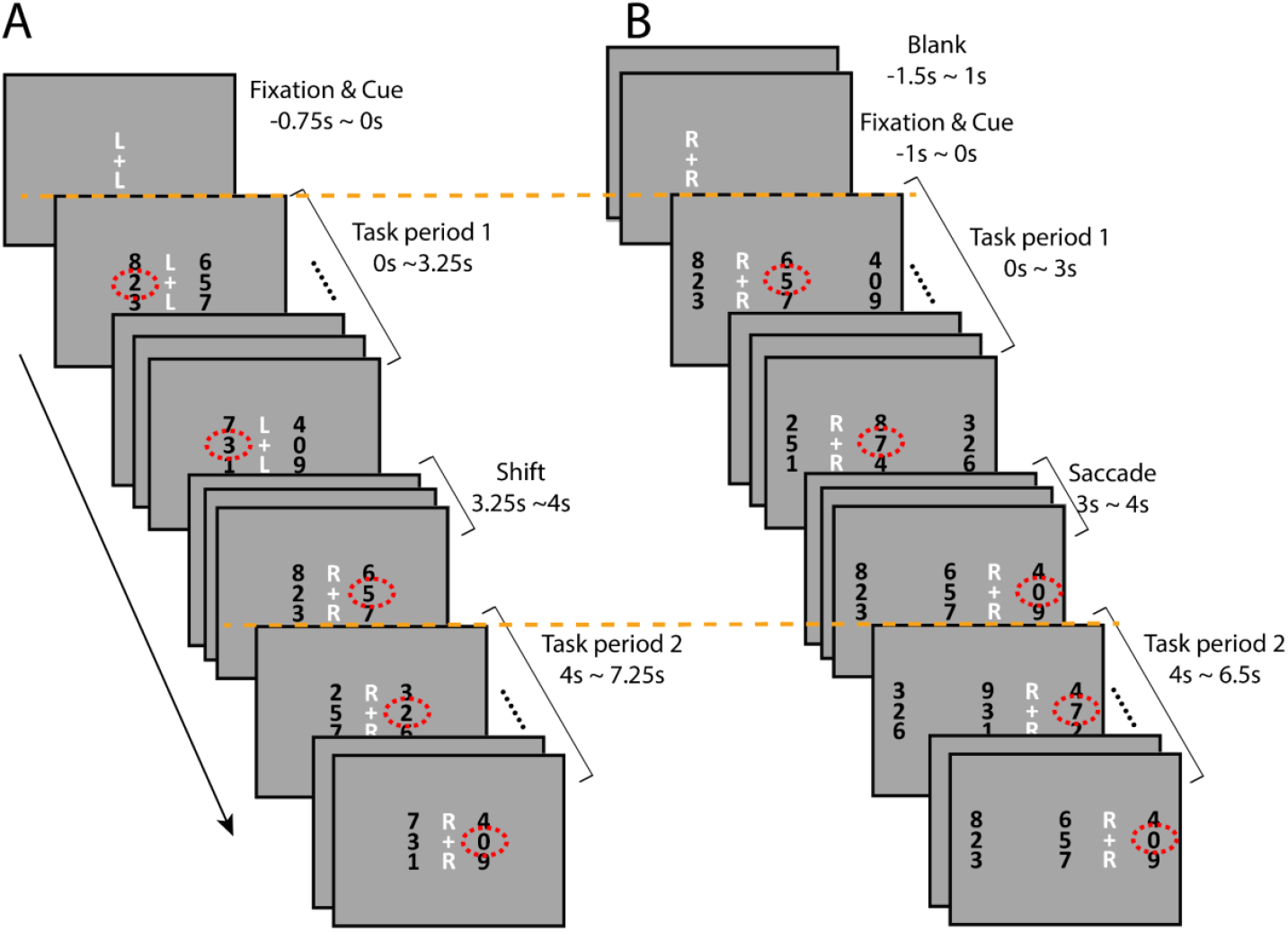
Paradigms of the Eyes-fixed and Eyes-move tasks. (A) an example of a shift-attention trial, from left of fixation to right of fixation; the letter cue “L” and “R” above and below the fixation indicates “left” and “right”. (B) an example of a retinotopic-attention trial, attention right of fixation across the saccade; the letter cue “L”, “R”, and “C” (not shown in this example) above and below fixation indicates “left of fixation”, “right of fixation”, and “center of screen”, respectively. Red dotted circles (not shown in the actual experiment) indicate the digit stream that participants should attend to according to the letter cue. Time 0s is taken as the onset of each trial, and orange dotted lines are to show that the onsets of task period 1 and 2 were synced with scanner pulse in both Eyes-fixed and Eyes-move tasks.

In the Eyes-fixed task (Figure 1A), subjects fixated their eyes at the fixation cross at the screen center. A letter cue appeared above and below the fixation to indicate the location to be covertly attended (L for left of fixation, R for right of fixation). The stimuli were rapid serial visual presentation (RSVP) streams of random digits (each frame of digits presented for 250ms without gap). Two columns of RSVP streams were located 2.5° to the left and right of the fixation cross, respectively. In each column, the middle stream was the target stream and the upper and lower streams were the flanker streams. Subjects were instructed to attend to the cued side and press the button when they saw a target (the number “5”) in the target stream.

Each trial lasted 8 seconds. The fixation and letter cue alone were presented for 0.75s before the onset of the RSVP streams. On half of the trials, the letter cue changed (e.g., from L to R) midway through the trial (always 3.25s after the onset of the RSVP streams), cueing participants to shift their covert attention to the other side and monitor for the target digit on the new side. Each trial can thus be thought of as containing two task periods, each lasting for 3.25s, separated by a 0.75s gap for the potential shift. (The RSVP streams continued during this potential shift period, but the target number “5” was inhibited.) The task was programmed so that the onset of the first task period was always synced with the scanner pulse (time 0 for each trial). The attended locations of the two periods could either be the same (“Hold-L” and “Hold-R” conditions) or different (“shift-LR” and “shift-RL” conditions), as shown in Figure 2. The four trial types were randomly intermixed in each Eyes-fixed run so that participants could not predict the conditions before each trial.

**Figure 2.**
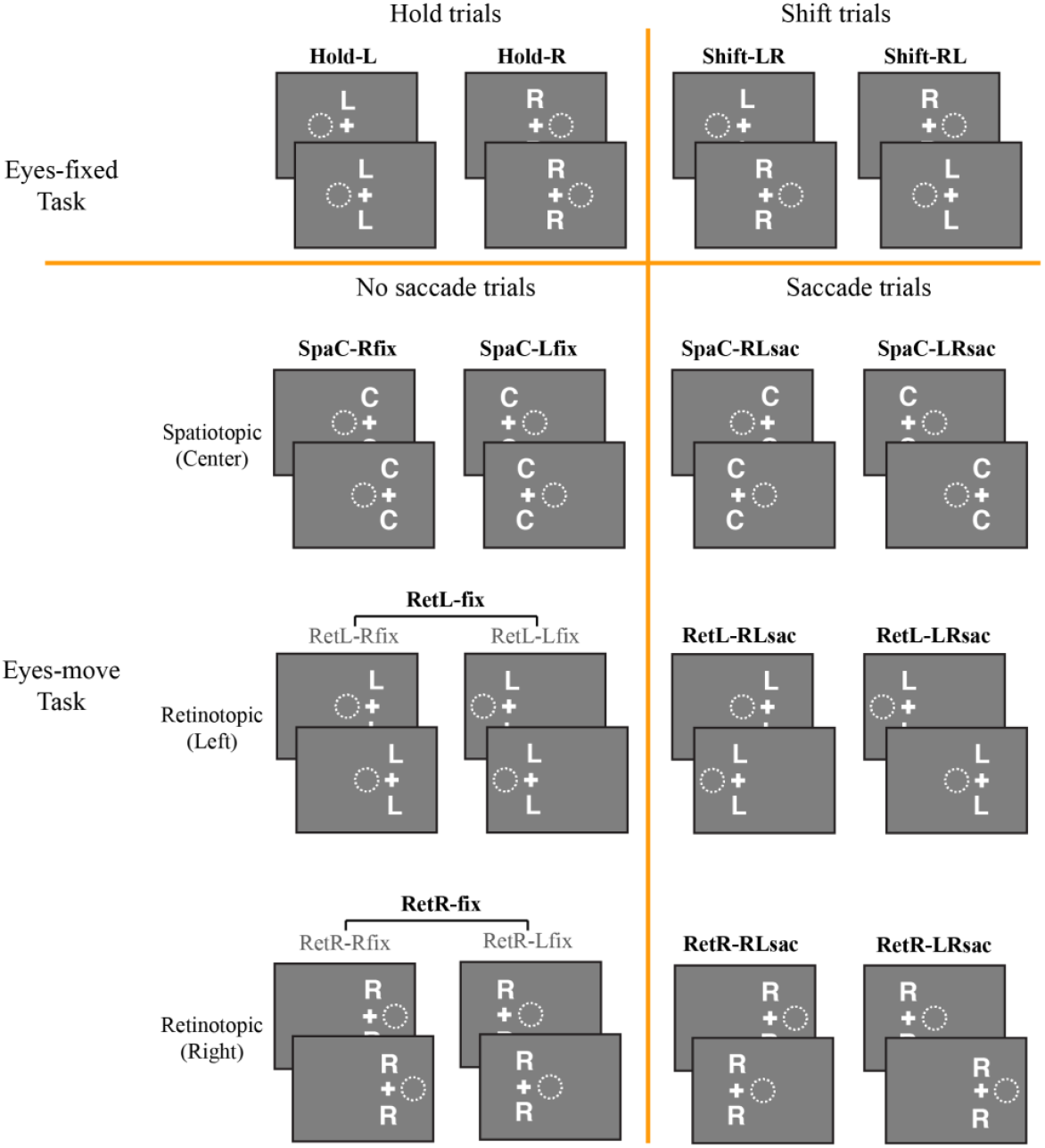
Diagrams of all conditions. Each condition was separated into the first half (before shift/saccade) and the second half (after shift/saccade), shown as the top and bottom panel for each condition. White crosses indicate the fixation location, and white dotted circles indicate the attention location on the screen, corresponding to the letter cues above and below the fixation. Note that in our analyses, we did not separate the left and right fixation for retinotopic no-saccade conditions; that is, only the bolded conditions were included in the GLMs.

The RSVP streams were composed of digits; the digit “5” was reserved as the target; other digits were presented randomly in a trial. In the RSVP, for every frame of 250 ms, there was a 1/3 chance that the target “5” would appear on the screen in one of middle (target) streams (when it appeared, it was randomly assigned to one of the target streams, and “5” never appeared in the flanker streams). The target presentation was temporally restricted so that two targets could not appear sequentially within 1s, no matter whether it appeared in the cued or uncued stream.

Stimuli in the Eyes-move task were similar, except that instead of fixating at the screen center, the fixation cross could appear at one of two potential fixation locations at the start of each trial, 2.5° to the left and right of the screen center, and there were three columns of RSVP streams, located at the far left, center, and far right of the screen, each centered 2.5° away from the nearest fixation location (Figure 1B). On half of the trials, the fixation cross remained in the same position for the entire trial (no-saccade trials); on the other half of trials, the fixation cross jumped to the other fixation location halfway through the trial (saccade trials). Subjects were instructed to fixate their eyes on the fixation cross and saccade to the new location whenever it moved.

Each Eyes-move run was subdivided into 4 mini-blocks (8 trials each). Two of these blocks contained the spatiotopic reference frame condition, where subjects were instructed to attend to the central RSVP stream regardless of where their eyes were. This condition was cued at the beginning of the mini-block as “attend screen center”, and the letter cue “C” appeared above and below the current fixation to remind subjects of the attended location. The other two mini-blocks contained the retinotopic reference frame conditions, where subjects were instructed to attend to an RSVP stream defined relative to fixation, i.e., “left of the cross” or “right of the cross”. These conditions were cued as such at the beginning of the mini-block, and with the letters “L” and “R”, respectively, during the trial. The order of these four mini-blocks was randomized in each run. Participants always knew which reference frame condition they were in, but they could not predict either the initial fixation location or whether they would have to make a saccade or not on each trial.

Each trial in the Eyes-move task also lasted 8 seconds. As in the Eyes-fixed task, the scanner pulse was always synced with the onset of the first task period (time 0); the rest of the trial was designed so that the time-course data would be as comparable as possible between Eyes-fixed and Eyes-move tasks. The initial fixation and letter cue alone appeared 1s before the start of the trial (onset of the RSVP streams). The first task period lasted 3s and the second 2.5s, separated by a 1s gap for a potential saccade. (The RSVP streams continued during this potential saccade period, but the target number “5” was inhibited.) There were another 0.5s of blank gap after the second task period before the next trial began.

A summary of all conditions in the Eyes-move task is listed in Figure 2. The conditions were coded based on reference frame, attended location, and fixation location or saccade direction. For example, in spatiotopic blocks, no-saccade trials were coded as SpaC-Rfix (spatiotopic reference frame, attend center stream, fixation on the right cross) and SpaC-Lfix, and saccade trials were coded as SpaC-RLsac (spatiotopic reference frame, attend center stream, saccade from right to left cross) and SpaC-LRsac. In retinotopic blocks, no-saccade trials were coded as RetL-Rfix (retinotopic reference frame, attend stream left of fixation, fixation on the right cross), RetL-Lfix, RetR-Rfix, and RetR-Lfix; however, although our design included both left and right fixation location trials, we aggregated them into RetL-fix and RetR-fix to simplify our analyses. This is because the aggregated conditions did not involve a visual field difference, and any effect coming from pure fixation location difference is beyond the main scope of this study. Retinotopic saccade trials were coded as RetL-RLsac (retinotopic reference frame, attend stream left of fixation, saccade from right to left cross), RetL-LRsac, RetR-RL-sac, and RetR-LRsac. These conditions are all illustrated in Figure 2. In sum, our main MVPA analyses included a total of 10 task conditions. (We also conducted a descriptive univariate analysis with different numbers of conditions; see Results section for details.)

In both Eyes-fixed runs and Eyes-move runs, trial onset times were jittered, with inter-trial intervals (ITIs) of 0s, 2s, and 4s (50%, 35%, and 15% of trials, respectively), in a fast-event related fashion. An additional mini-block (16s) of blank baseline was put in the beginning, middle and end of each run, respectively, where participants were instructed to keep fixated at the fixation cross. Participants completed 4 runs of Eyes-fixed task and 8 runs of Eyes-move task. In addition, they also completed 2 to 4 runs of the standard retinotopic mapping task (see details in the ROI section below).

All stimuli were generated with the Psychtoolbox (Brainard, 1997) in Matlab (MathWorks). Stimuli were displayed with a 3-chips DLP projector onto a screen in the rear of the scanner (resolution 1280×1024 at 60Hz). Participants viewed from a distance of 74cm via a mirror above attached to the head coil.

### Eye Tracking

Eye positions were recorded throughout the experiment when the calibration was reliable, using an MRI-compatible Eyelink remote eye-tracker at 500 Hz. Eye position data were used to ensure the participants kept their eyes on the fixation point and made eye movements following the fixation change. When eye position data were not available, the experimenters observed the subjects’ eye through the camera and made sure that the participants were making eye movements as intended.

### fMRI acquisition

This study was done at [anonymous location] with a Siemens Prisma 3T MRI scanner using a 32-channel phase array receiver head coil. Functional data were acquired using a T2-weighted gradient-echo sequence (TR=2000ms, TE=28ms, flip angle 71°). The slice coverage was oriented about 45° away from the AC-PC plane and placed to prioritize full coverage of occipital and parietal lobes, and then maximize coverage of temporal and frontal lobes (33 slices, 2×2×2mm voxel, 10% gap). We also collected a high-resolution MPRAGE anatomical scan at 1mm3 resolution for each participant. Each participant was scanned in one two-hour session.

### fMRI preprocessing

The fMRI data were preprocessed with Brain Voyager QX (Brain Innovation). All functional data were corrected for slice acquisition time and head motion, temporally filtered. Runs with abrupt motion greater than 1mm were discarded from later analyses, and the motion correction parameters were logged and input as nuisance variables into the GLM. Spatial smoothing of 4mm FWHM was performed on the preprocessed data for univariate analyses, but not for multivoxel pattern analysis (MVPA). Data of each participant were normalized into Talairach space (Talairach & Tournoux, 1988). We used FreeSurfer to segment the white matter / gray matter boundaries from each participant’s anatomical scan, and imported the images into BrainVoyager for flattening. We extracted each participant’s cortical surface for each hemisphere in Talairach space, and inflated and flattened them into cortical surface space for retinotopic mapping. Other analyses were performed on volume space only.

### Regions of Interest

We defined four a priori regions of interest (ROIs). Two of these ROIs were our theoretical regions of interest designed to look at attentional representations: bilateral area V4 (considered strongly modulated by attention: McAdams & Maunsell,2000;), and an attention shift network (functionally defined as described below). We also defined two comparison ROIs to capture visual activation (bilateral area V1) and deactivation / default mode (functionally-defined task negative network).

The attention shift network was functionally defined based on the group-level shift > hold univariate attention contrast in the Eyes-fixed task. For this contrast, we used a whole-brain multi-subject general linear model (GLM) in the Eyes-fixed task with 5 regressors (blank baseline plus the 4 Eyes-fixed conditions) and 6 nuisance regressors from the motion correction processing, with a canonical hemodynamic response function, to calculate beta weights of each condition for each voxel. We then projected the contrasts of shift conditions vs hold conditions onto volume maps. All volume maps were corrected for cluster threshold at α=0.05 level, using the BrainVoyager plugin “Cluster-level Statistical Threshold Estimator”, after which all significant voxel clusters were picked as the corresponding functional network. The task negative network was defined in a similar way, based on the group-level baseline > task contrast in the Eyes-fixed task, where task included all 4 Eyes-fixed task conditions.

The attention shift and task negative networks are shown in Figure 3 and Table 1. The attention shift network includes inferior parietal lobule (IPL) and temporal gyri, consistent with areas previously found in the literature (Beauchamp et al., 2001; Corbetta et al., 1998; de Haan et al., 2008; Yantis et al., 2002). The task negative network includes a posterior medial portion of parietal cortex, often considered part of the default mode network (Greicius et al., 2003). Due to limited frontal coverage in our scanning protocol, our data only captured more posterior regions.

**Figure 3.**
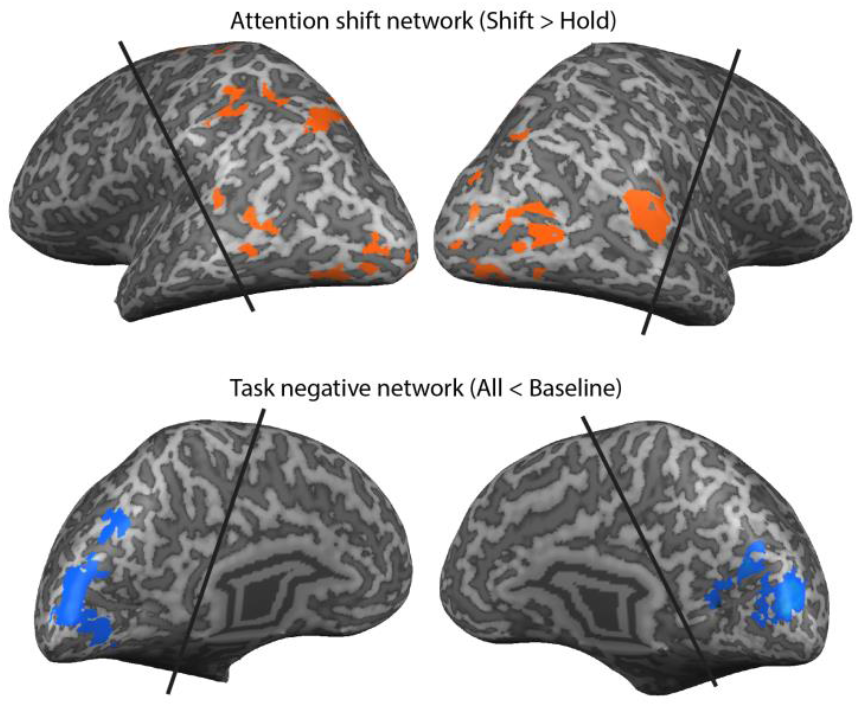
The attention shift network (above) and the task negative network (below). Result is based on group-level GLM contrasts on the volume space, after being cluster-threshold corrected at p<.05. The volume maps were projected onto an inflated brain only for visualization purpose. The black lines demonstrate the approximate coverage (slightly different for each subject).

**Table 1.**
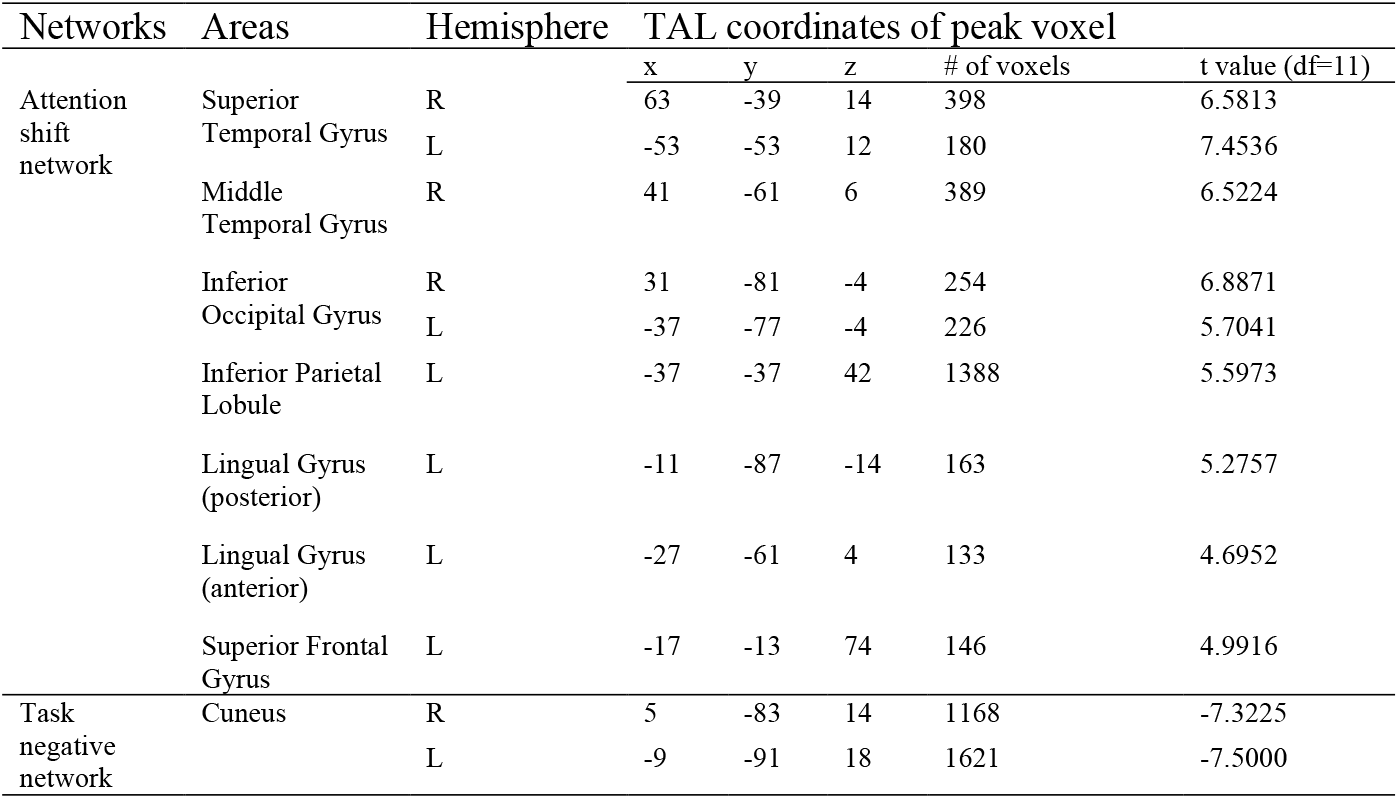
Description of clusters in the attention shift network and the task negative network, including Talairach coordinates of the peak voxel, number of voxels, and t values.

We used a standard retinotopic mapping localizer (Sereno et al., 1995) to define visual ROIs (V1 and V4) for each participant. In the localizer task, a rotating wedge with high-contrast radial checkerboard patterns was presented on the screen and flickered at 4 Hz. The 60° wedge stimulus covered eccentricity from 1.6° to 16° and was rotated either clockwise or counter-clockwise for 7 cycles with a period of 24 s per cycle. Participants were instructed to fixate at the center fixation of the screen, and press the button every time when the fixation dot changed color from dark grey to light grey. A pair of clockwise and counterclockwise runs were combined in the analyses. One or two pairs of runs (i.e., 2 to 4 runs) were obtained for each participant. After preprocessing, the brain data were analyzed in custom Matlab code and projected onto the flattened brains as surface maps in Brain Voyager. Bilateral V1 and V4 boundaries were defined based on these surface maps. We then used the task > baseline contrast from the Eyes-fixed runs to further constrain the retinotopic ROIs to regions visually activated by this task.

In addition to these four a priori regions of interest, we also defined a post-hoc network for exploratory analyses, the “retinotopic-hold” network, based on the cross-task similarity searchlight results (see details below), corrected for cluster threshold in the same way as above. ROI results for this post-hoc network are presented for descriptive purposes only, as the datasets used to define and analyze were not fully independent.

### Multivoxel pattern analyses (MVPA)

For all MVPA analyses below, we imported corresponding GLM data to Matlab with BrainVoyager’s BVQXtools Matlab toolbox, and all subsequent analyses were done using custom Matlab code.

#### 1) Correlation-based MVPA

We performed MVPA using the split-half correlation-based method (Haxby et al., 2001) for each participant and each ROI/network. We obtained GLMs for odd runs and even runs separately for each participant; each GLM had 5 regressors for the Eyes-fixed task (blank baseline plus the 4 Eyes-fixed conditions) and 11 regressors for the Eyes-move task (blank baseline plus the 10 Eyes-move conditions from Figure 2), as well as 6 nuisance regressors from the motion correction processing. For the following analyses, we focused on non-baseline conditions. For each GLM, we normalized the voxel data (beta weights) by subtracting the mean response across all non-baseline conditions from the response of each individual condition, for each voxel. The response patterns (voxel-wise beta weights after de-meaning) for each condition in the even runs were then correlated with the patterns for each condition in the odd runs, generating a correlation matrix for each task. The correlation coefficients were transformed into z-scores using Fisher’s r-to-z transform.

We then calculated the following types of information based on the correlation matrix. In the Eyes-fixed task: information about shift execution (holding vs shifting attention), hold attention location (holding left vs holding right), and shift direction (shifting leftward vs shifting rightward). In the Eyes-move task: information about saccade execution (saccade vs no saccade), saccade direction (saccade leftward vs saccade rightward), and reference frame (attend retinotopic task vs attend spatiotopic task). Specifically, we picked out cells in the matrix that reflected the same type of information (“within-category” correlations, e.g., holding attention correlated with holding attention), and cells that reflected the different type of information (“between-category” correlations, e.g., holding attention correlated with shifting attention). The information index was then calculated by subtracting the mean correlation values of “different” cells from those of “the same” cells. A significantly-positive information index value would indicate that there is some amount of information of this type in the ROI.

#### 2) Multivoxel Pattern Time Course (MVPTC) analyses

The first step of analyses described above used regular whole-trial GLMs, which modeled the whole 8 sec (4 TR) trial as a single event. However, since trials contained a potential attention shift or saccade halfway through, the initial analysis might fail to capture some dynamic brain representations. Thus, we also performed timecourse analyses using finite impulse response (FIR) GLM analyses with 10 timepoints, on the same conditions as above. Timepoint zero (TP0) corresponds to the start of the first task period in each trial (i.e., the onset of RSVP stimuli). We fed those FIR GLMs into MVPA analyses (i.e., MVPTC, modified from Chiu, Esterman, Gmeindl, & Yantis, 2012). Taking each time point as a separate dataset, we performed similar analyses as above to calculate the information indices. The result figures show all 10 TPs in the FIR, but our statistical analyses focus on three TPs that capture critical time periods in the trial, accounting for BOLD signal lag: TP3 (before the shift/saccade happened), TP4 (capturing the shift/saccade), TP5 (after the shift/saccade). It is also important to clarify that at the behavioral time period corresponding to BOLD signals at TP3, participants did not know yet whether there would be an attentional shift or not (in eyes-fixed task), or a saccade or not (in Eyes-move task), because the trials were intermixed; however, it was predictable that if there would be a shift/saccade, what direction the shift/saccade would be, based on the attention location or the eye location within the first half of a trial.

#### 3) Cross-task pattern similarity analysis

To directly compare the similarity *between* the brain activity patterns of covert attention during Eyes-fixed and Eyes-move tasks, we also performed a cross-task pattern similarity analysis for both whole-trial and time-course beta weights. Because the Eyes-fixed and Eyes-move tasks were performed in separate runs, we used GLMs of all runs instead of split-half to increase power; that is, we took Eyes-fixed runs and Eyes-move runs as the two datasets for the correlation analysis. After de-meaning the voxel-wise responses in the same way as above, we calculated the z-scored correlation matrix comparing each condition in the Eyes-fixed task to each saccade condition in the Eyes-move task. We then calculated the pattern similarity between holding, shifting, retinotopic, and spatiotopic attention by averaging the z-scored correlation coefficients of corresponding cells in the matrix. The similarity data were submitted to a 2 (Eyes-move conditions: retinotopic & spatiotopic) by 2 (similarity to Eyes-fixed conditions: similarity-to-hold & similarity-to-shift) ANOVA, with four cross-task similarities total (i.e., four pairs of correlations). In this ANOVA analysis, a main effect of similarity to Eyes-fixed conditions would indicate that both retinotopic and spatiotopic attention are represented more similarly to hold (or shift) attention than shift (or hold); an interaction would indicate relatively greater similarity between retinotopic and holding attention & between spatiotopic and shifting attention (or relatively greater similarity between spatiotopic and holding attention & between retinotopic and shifting attention).

#### 4) Whole-brain searchlight on cross-task pattern similarity analysis

Finally, we performed MVPA searchlight analyses (Kriegeskorte et al., 2006) to search across the entire slice coverage, for clusters that might show patterns of interest outside our a priori ROIs. The approach is similar to what is described above; instead of taking a priori ROIs, we searched through individual brains iteratively with a “moving” ROI, defined as a sphere of radius 3 voxels. On each iteration, MVPTC analyses were performed as described above on each ROI sphere, and z-scored correlation values were assigned to the center voxel of this ROI sphere to form z-maps for each subject. Specifically, we focused on the interaction term in the 2 (Eyes-move conditions: retinotopic & spatiotopic) by 2 (similarity to Eyes-fixed conditions: similarity-to-hold & similarity-to-shift) ANOVA and focused only on TP4, which theoretically captured the timepoint at shift/saccade. In practice, we first generated 4 searchlight maps for each individual subject, indexing each pair of correlations: retinotopic-to-hold, spatiotopic-to-hold, retinotopic-to-shift, and spatiotopic-to-shift. We calculated the difference (subtraction) between the first and the second maps, as well as the difference between the third and fourth similarity maps, before calculating the final difference of differences (interaction) map for each subject. The interaction maps for each individual were then spatially smoothed with a 4 mm FWHM kernel to facilitate a group analysis. The group t-value map was constructed using two-tailed t-tests comparing the interaction values for each voxel against zero, correcting for cluster threshold in the same way as above. A positive t-value for a given voxel indicates that retinotopic attention is represented more similar to holding attention, and spatiotopic more similar to shifting attention (i.e., the “retinotopic-hold / spatiotopic-shift” pattern); in contrast, a negative t-value indicates that retinotopic attention is represented more similar to shifting attention and spatiotopic more similar to holding attention (i.e., the “spatiotopic-hold / retinotopic-shift” pattern).

## Results

### Behavior results

To evaluate participants’ behavioral performance, we defined hits as correctly pressing a button within 1 sec in response to a “5” target at the attended location and false alarms as incorrectly pressing a button when there was no “5” target within 1 sec at the attended location. We calculated the hit rate by dividing the total number of hits in each trial by the total number of targets at the attended location (trials with 0 targets were omitted). We also calculated the false alarm rate by dividing the total number of false alarms in each trial by the total number of frames when there was no target presented in the attended RSVP stream. D-prime was calculated by subtracting z-scored false alarm rates from z-scored hit rates.

Due to a coding mistake for data logging, two subjects did not have reliable behavioral responses logged and were excluded from the analyses of behavioral performance. For the remaining 10 subjects, the mean hit rate in Eyes-fixed task was 66.17% (±5.07% standard deviation), and the mean false alarm rate was 0.52% (±0.14% standard deviation); in Eyes-move task, the mean hit rate was 65.67% (±5.70%) and the mean false alarm rate was 0.50% (±0.18%). These two tasks were designed to be hard to make sure that participants maintained attention on the cued location, so it is reasonable that participants’ performance was not at ceiling. The d-prime measurements in both tasks were well above zero, *t*’s≥15.239, *p*’s≤.001, Cohen’s *d*’s≥4.819, and there was no significant difference between the two tasks, *t*(9)=0.217, *p*=.833, Cohen’s *d*=0.069. In addition, there were no significant differences of d-prime between hold and shift attention in Eyes-fixed task, between saccade and no saccade trials in Eyes-move task, and between spatiotopic and retinotopic attention, all *t*’s≤2.083, *p*’s≥.067, Cohen’s *d*’s≤0.659.

### Univariate results

To give a general view of how the brain activity looks like for each condition, Figure 4 plots the percent signal change in the time course as well as the univariate beta weights for each a priori ROI. To better illustrate, we recoded the conditions to plot them according to whether the attended side was ipsilateral/contralateral relative to the ROIs in each hemisphere, and further collapsed across the RL and LR saccade directions in retinotopic saccade trials (that is, only 8 conditions were shown in results). To make it comparable for each condition, we subtracted the percent signal change or beta weights of fixation baseline from all other conditions, in both Eyes-fixed and Eyes-move task. As shown in Figure 4, there was a separation in the attention shift network between holding and shifting attention around TP4, as well as a clear pattern of contralateral attentional modulation in V4, but no major differences in other ROIs/networks between conditions.

**Figure 4.**
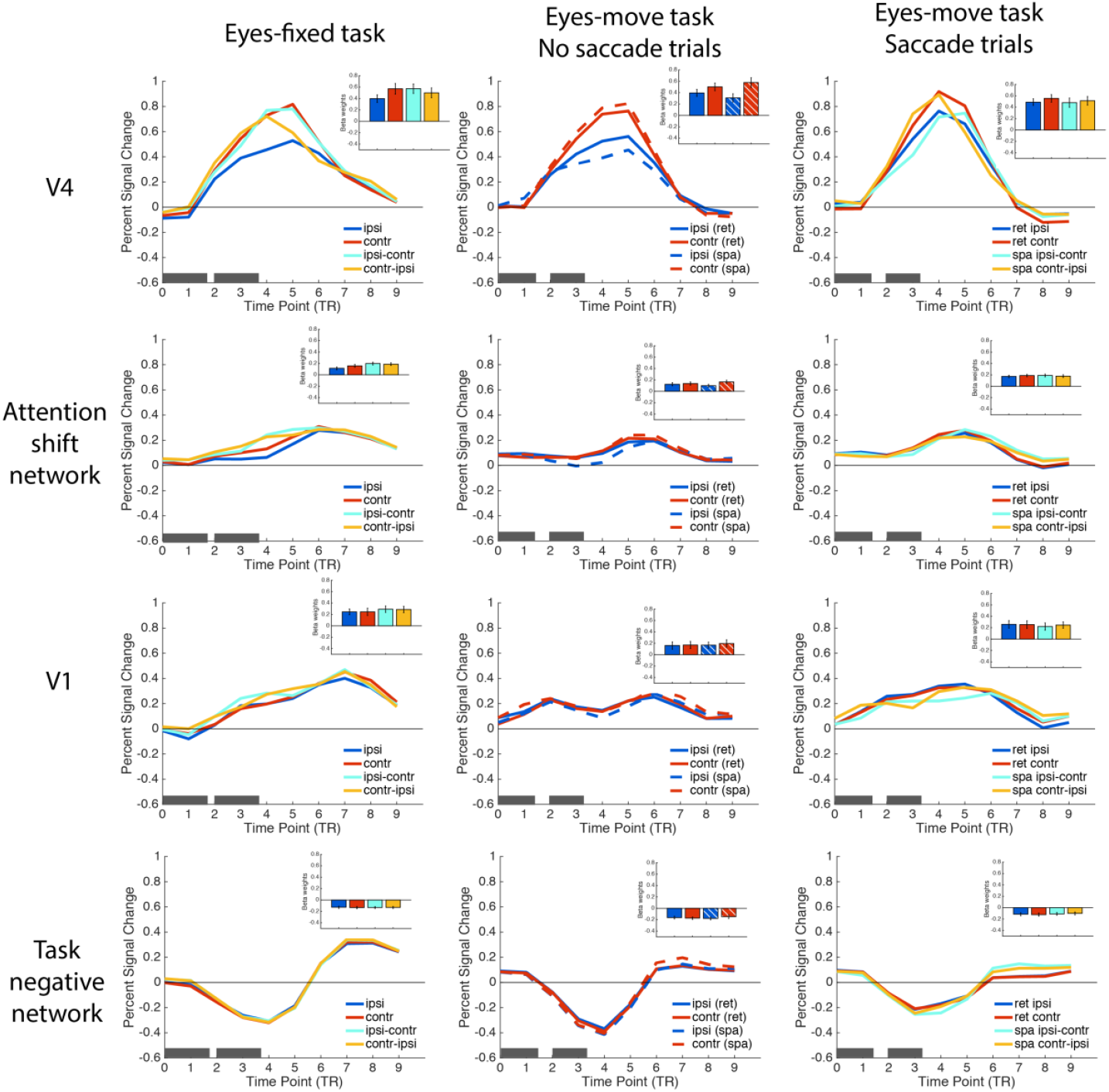
FIR results of Eyes-fixed task (A), Eyes-move task with no saccade trials (B), and Eyes-move task with saccade trials (C). The pair of gray boxes along the x-axis in each plot indicates the time duration of the two task periods in the trial. Inset bar plots show the whole-trial beta weights for each condition in each ROI/network, color-coded in the same way as the corresponding FIR plots.

### MVPA of shifting vs holding attention (Eyes-fixed)

For the Eyes-fixed task, we asked whether we could decode from the brain patterns information about shift execution (holding vs shifting attention trials), about hold attention location (attending left vs right stream on hold trials), and about shift direction (shift left-right vs shift right-left trials) (Figure 5A). From each of our a priori ROIs/networks, we conducted correlation-based MVPA on the whole-trial GLM beta weights (Figure 5B). We also asked how these three types of information develop over the time course of the trials (MVPTC), by using beta weights from the FIR GLMs (Figure 5C). As explained in the methods section, we particularly focused on the BOLD signals at TP3, TP4, and TP5, which corresponded to the three critical behavioral time periods, including before the shift/saccade happened, around the shift/saccade, and after the shift/saccade was done. Statistics of all one-sample t-tests on the information indices are listed in Table 2. For whole-trial MVPA, we corrected for multiple comparisons across the four ROIs/networks using Holm–Bonferroni correction; for the MVPTC analysis in each ROI/network, we corrected for multiple comparisons across the three TPs. The results for three types of information are summarized separately below.

**Figure 5.**
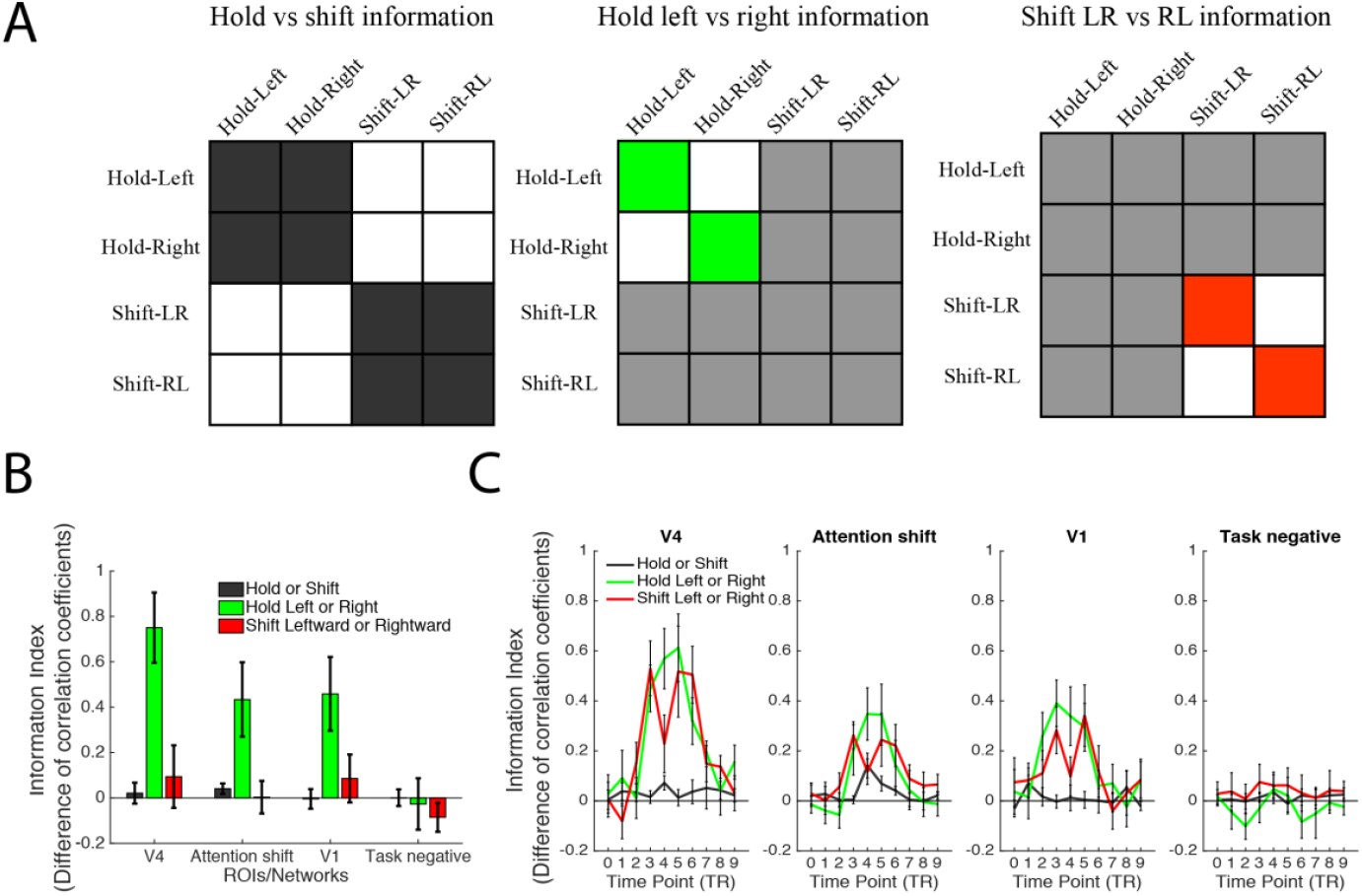
MVPA and MVPTC analyses and results of Eyes-fixed tasks. (A) Hypothetical matrices for hold vs shift, hold left vs right, and shift LR vs RL information. Cells colored in dark grey, green and red are the within-group correlations, and white cells are the between-group correlations. Light grey cells are not used in the corresponding analysis. The index value of each type of information is calculated by subtracting the z-scored between-group correlation coefficients from the z-scored within-group correlation coefficients. (B) The index value of each type of information in each ROI/network. (C) The index value of each type of information at 10 time points, in each ROI/network. Error bars represent SEM.

**Table 2.** Statistical tests of information indices in each ROI/network, separately for whole-trial analyses and time points of interest in the time-course analyses. N=12.

|  | V4 | Attention shift network | V1 | Task negative network |
| --- | --- | --- | --- | --- |
| Hold<br>or<br>Shift | $t(11)=0.451, p=.661, d=0.130$<br>TP3: $t(11)=0.608, p=.555, d=0.176$<br>TP4: $t(11)=2.507, p=.029, d=0.724^*$<br>TP5: $t(11)=0.394, p=.701, d=0.114$ | $t(11)=1.755, p=.107, d=0.507$<br>TP3: $t(11)=0.278, p=.787, d=0.080$<br>TP4: $t(11)=2.853, p=.016, d=0.823^{**}$<br>TP5: $t(11)=2.316, p=.041, d=0.668^*$ | $t(11)=-0.102, p=.920, d=-0.030$<br>TP3: $t(11)=-0.104, p=.919, d=-0.030$<br>TP4: $t(11)=0.236, p=.818, d=0.068$<br>TP5: $t(11)=0.127, p=.902, d=0.037$ | $t(11)=0.016, p=.987, d=0.005$<br>TP3: $t(11)=0.751, p=.469, d=0.217$<br>TP4: $t(11)=1.451, p=.175, d=0.419$<br>TP5: $t(11)=-0.339, p=.741, d=-0.098$ |
| Hold L<br>or<br>Hold R | $t(11)=4.843, p<.001, d=1.380^{**}$<br>TP3: $t(11)=4.818, p<.001, d=1.391^{**}$<br>TP4: $t(11)=4.709, p<.001, d=1.359^{**}$<br>TP5: $t(11)=4.521, p<.001, d=1.305^{**}$ | $t(11)=2.645, p=.023, d=0.764^{**}$<br>TP3: $t(11)=2.025, p=.068, d=0.585$<br>TP4: $t(11)=3.326, p=.007, d=0.960^{**}$<br>TP5: $t(11)=2.834, p=.016, d=0.818^{**}$ | $t(11)=2.821, p=.017, d=0.815^{**}$<br>TP3: $t(11)=4.224, p=.001, d=1.219^{**}$<br>TP4: $t(11)=2.880, p=.015, d=0.831^{**}$<br>TP5: $t(11)=3.071, p=.011, d=0.887^{**}$ | $t(11)=-0.237, p=.817, d=-0.068$<br>TP3: $t(11)=-0.498, p=.628, d=-0.144$<br>TP4: $t(11)=0.813, p=.434, d=0.235$<br>TP5: $t(11)=0.287, p=.780, d=0.083$ |
| Shift leftward<br>or<br>rightward | $t(11)=0.682, p=.510, d=0.197$<br>TP3: $t(11)=4.840, p<.001, d=1.397^{**}$<br>TP4: $t(11)=1.975, p=.074, d=0.570$<br>TP5: $t(11)=2.839, p=.016, d=0.820^{**}$ | $t(11)=0.040, p=.969, d=0.012$<br>TP3: $t(11)=4.903, p<.001, d=1.415^{**}$<br>TP4: $t(11)=2.273, p=.044, d=0.656^{**}$<br>TP5: $t(11)=2.310, p=.041, d=0.667^*$ | $t(11)=0.808, p=.437, d=0.233$<br>TP3: $t(11)=3.013, p=.012, d=0.870^{**}$<br>TP4: $t(11)=1.247, p=.238, d=0.360$<br>TP5: $t(11)=2.772, p=.018, d=0.800^{**}$ | $t(11)=-1.334, p=.209, d=-0.385$<br>TP3: $t(11)=1.059, p=.312, d=0.306$<br>TP4: $t(11)=1.153, p=.273, d=0.333$<br>TP5: $t(11)=0.886, p=.394, d=0.256$ |
| Saccade<br>or<br>no saccade | $t(11)=1.452, p=.175, d=0.419$<br>TP3: $t(11)=2.056, p=.064, d=0.594$<br>TP4: $t(11)=3.305, p=.007, d=0.954^{**}$<br>TP5: $t(11)=2.014, p=.069, d=0.581$ | $t(11)=4.432, p=.001, d=1.279^{**}$<br>TP3: $t(11)=2.598, p=.025, d=0.750^{**}$<br>TP4: $t(11)=2.956, p=.013, d=0.853^{**}$<br>TP5: $t(11)=7.249, p<.001, d=2.093^{**}$ | $t(11)=1.522, p=.156, d=0.439$<br>TP3: $t(11)=0.819, p=.430, d=0.236$<br>TP4: $t(11)=2.552, p=.027, d=0.737^*$<br>TP5: $t(11)=0.977, p=.350, d=0.282$ | $t(11)=2.527, p=.028, d=0.730^*$<br>TP3: $t(11)=2.517, p=.029, d=0.727^{**}$<br>TP4: $t(11)=2.888, p=.015, d=0.834^{**}$<br>TP5: $t(11)=4.115, p=.002, d=1.188^{**}$ |
| Saccade<br>leftward<br>or<br>rightward | $t(11)=2.730, p=.020, d=0.788^*$<br>TP3: $t(11)=4.113, p=.002, d=1.187^{**}$<br>TP4: $t(11)=7.401, p<.001, d=2.136^{**}$<br>TP5: $t(11)=3.370, p=.006, d=0.973^{**}$ | $t(11)=1.771, p=.104, d=0.511$<br>TP3: $t(11)=4.420, p=.001, d=1.276^{**}$<br>TP4: $t(11)=1.775, p=.104, d=0.512$<br>TP5: $t(11)=3.615, p=.004, d=1.044^{**}$ | $t(11)=0.021, p=.984, d=0.006$<br>TP3: $t(11)=1.329, p=.211, d=0.384$<br>TP4: $t(11)=0.072, p=.944, d=0.021$<br>TP5: $t(11)=1.219, p=.248, d=0.352$ | $t(11)=2.602, p=.025, d=0.751^*$<br>TP3: $t(11)=3.699, p=.004, d=1.068^{**}$<br>TP4: $t(11)=3.228, p=.008, d=0.932^{**}$<br>TP5: $t(11)=3.484, p=.005, d=1.006^{**}$ |
| Retinotopic<br>or<br>spatiotopic | $t(11)=0.504, p=.625, d=0.145$<br>TP3: $t(11)=0.067, p=.948, d=0.019$<br>TP4: $t(11)=0.101, p=.921, d=0.029$<br>TP5: $t(11)=0.819, p=.430, d=0.237$ | $t(11)=0.074, p=.943, d=0.021$<br>TP3: $t(11)=0.816, p=.432, d=0.236$<br>TP4: $t(11)=1.295, p=.222, d=0.374$<br>TP5: $t(11)=1.095, p=.297, d=0.316$ | $t(11)=1.225, p=.246, d=0.354$<br>TP3: $t(11)=0.825, p=.427, d=0.238$<br>TP4: $t(11)=3.498, p=.005, d=1.010^{**}$<br>TP5: $t(11)=1.138, p=.279, d=0.329$ | $t(11)=0.268, p=.794, d=0.077$<br>TP3: $t(11)=0.662, p=.522, d=0.1911$<br>TP4: $t(11)=2.367, p=.037, d=0.683^*$<br>TP5: $t(11)=0.779, p=.452, d=0.225$ |

\* indicate statistical significance at $p<.05$ \*\* indicate statistical significance at $p<.05$ (Holm-Bonferroni corrected for multiple post hoc comparisons, separately across ROIs/networks for whole-trial MVPA, and across three TPs for MVPTC)

For the information about shift execution (holding vs shifting attention trials), we did not find significant information with the whole trial MVPA analyses in the attention shift network, nor in other ROIs/networks. However, recall that trials were 8-seconds long, and the hold and shift trials were designed to be identical for the majority of the trial, except the transient shift occurring midway through the trial. Indeed, when analyzing the time course in the attention shift network, we did find significant information about shift execution at the critical TP4. There was a weak effect at TP5 that did not pass correction for multiple comparisons, and no significant information about shift execution for TP3, before the shift happened. The MVPTC analyses thus successfully captured a transient change in activity pattern around the time when the shifts happened, in the attention shift network^1^. In V4 we found information about shift execution at TP4 that did not pass correction for multiple comparisons. No significant information about shift execution was found in V1 or the task negative network for any timepoints.

For the information about which location was attended on hold trials (holding left vs right stream), we found significant information in the whole-trial MVPA, in both the attention shift network and V4. The information was also significant in V1, but not in the task negative network. MVPTC showed that this information was sustained for the duration of the trial and was significant at TP3, TP4 and TP5 in the attention shift network, V4, and V1, with the only exception at TP3 in the attention shift network. This is consistent with the behavioral task on these hold trials, in that participants maintained attention in one location throughout the entire trial, indicating that the information about the location attended was represented broadly in all of our visual and attention ROIs/networks, but not in the task negative network.

The analogous analysis for the shift attention trials asks if we can decode information about covert attention shift direction (shift left-right vs shift right-left trials). We did not find significant information in any of the ROIs/networks with whole-trial beta weights. The timecourse analyses may give some insight into why. Interestingly, the MVPTC took a different shape than for the previous analyses; here, instead of peaking at the critical TP4, the information was actually greater at TP3 and TP5 than at TP4 in most ROIs/networks (except for the task negative network, which contained no significant information at any of the 3 timepoints). In V1 and V4, the shift direction information was significant at TP3 and TP5 but not TP4. This bimodal pattern also existed in the attention shift network numerically, but all three TPs were significant. It should be noted that in our design, the direction of the shifting was perfectly confounded with the location participants attended to before and after the shift. Thus, the bimodal pattern may reflect a dynamic representation of which location was being attended in the first half of the trial (peaking at TP3), and then after the attention shift in the second half of the trial (peaking at TP5), rather than reflecting information about the shift direction itself.

### MVPA of attention maintained across saccades (Eyes-move)

For the Eyes-move task, we used a similar approach of whole-trial MVPA followed by MVPTC to ask whether the activity patterns in each ROI contained information about saccade execution (saccade vs no-saccade trials), and on saccade trials, about the saccade direction (leftward vs rightward saccade) and about reference frame (retinotopic vs spatiotopic attention) (Figure 6, statistics in Table 2).

For the information about saccade execution (saccade vs no-saccade trials), we found significant information in whole-trial MVPA analyses in the attention shift network. There was also a weak effect not passing correction in the task negative network. When looking at time-course analyses, we found that the information was represented significantly in V4 and the attention shift network at TP4, corresponding to the behavioral time period of saccade execution. In the attention shift network, this information was also significant at both TP3 and TP5. Post hoc t-tests comparing the information indices at TP3/TP5 to TP4 showed that the information at the critical TP4 was significantly greater than at TP3, *t*(11)=2.772, *p*=.018, Cohen’s *d*=0.800, but information at TP4 was only numerically larger than at TP5, *t*(11)=0.946, *p*=.364, Cohen’s *d*=0.273. It is possible that saccade preparation and saccade execution might have elongated the process and thus blurred the effect temporally in the attention shift network. This information was also significant at TP4 in the task negative network, but did not survive correction at TP4 in V1.

For the information about saccade direction (right-left saccade vs left-right saccade), we found weak information that did not pass correction with whole-trial MVPA, in both V4 and the task negative network, but not in the attention shift network or V1. In the MVPTC, the saccade direction information was significant in all three timepoints in V4, and at TP3 and TP5 in the attention shift network. This information was also significant in all three timepoints in the task negative network, and was not significant in any of these time points in V1. Some of the timecourses appeared to have a similar bimodal shape for information about saccade direction as above for covert attention shift direction, perhaps again driven by information about attended hemisphere over time (Extended data Figure 6-1). Interestingly, although both V4 and the attention shift network represented information on saccade execution and saccade direction information, V4 seems to have more information about saccade direction, whereas the attention shift network had more information about saccade execution.

Finally, we did not find reference frame information (retinotopic-attention vs spatiotopic-attention trials) in whole-trial MVPA analyses in any of the ROIs/networks. There was significant information about reference frame in V1 at TP4 and an effect that did not pass correction at TP4 in the task-negative network, but no timepoints were significant in V4 or the attention shift network. Thus, our primary attentionally-modulated ROIs did not appear to directly differentiate which reference frame participants were maintaining attention in, though as noted above, they contained information about which location was being attended at any given time, and whether saccades were being executed.

### Cross-task similarity analysis of covert attention at fixation and across saccades

The above results demonstrate that brain regions sensitive to attentional modulation (V4 and the attentional shift network) represent information about covert attention shifts and about saccade execution. Now the key question is, how do representations of covert attention during fixation compare to covert attention maintained across saccades? Depending on the reference frame, both spatiotopic and retinotopic attention could be thought of as “hold” or “shift” attention tasks: spatiotopic attention is maintained in the same location relative to the screen, but shifted relative to our eyes, whereas retinotopic attention is the opposite. Is one or both of these tasks represented more similarly to holding attention in some brain regions, and/or more similarly to shifting attention elsewhere in the brain? To answer these questions, we analyzed the pattern similarity between Eyes-fixed conditions and Eyes-move conditions (Figure 7A). Rather than calculate information indices, in this cross-task MVPA analysis we directly compare the representational similarity scores for each cross-task pair of conditions (i.e., similarity between retinotopic and hold, between spatiotopic and hold, between retinotopic and shift, between spatiotopic and shift). We submitted these data to a 2 (Eyes-move condition: retinotopic & spatiotopic) x 2 (Eyes-fixed comparison: similarity-to-hold & similarity-to-shift) repeated-measures ANOVA for each ROI/network at each critical timepoint, as well as for the whole trial.

**Figure 6.**
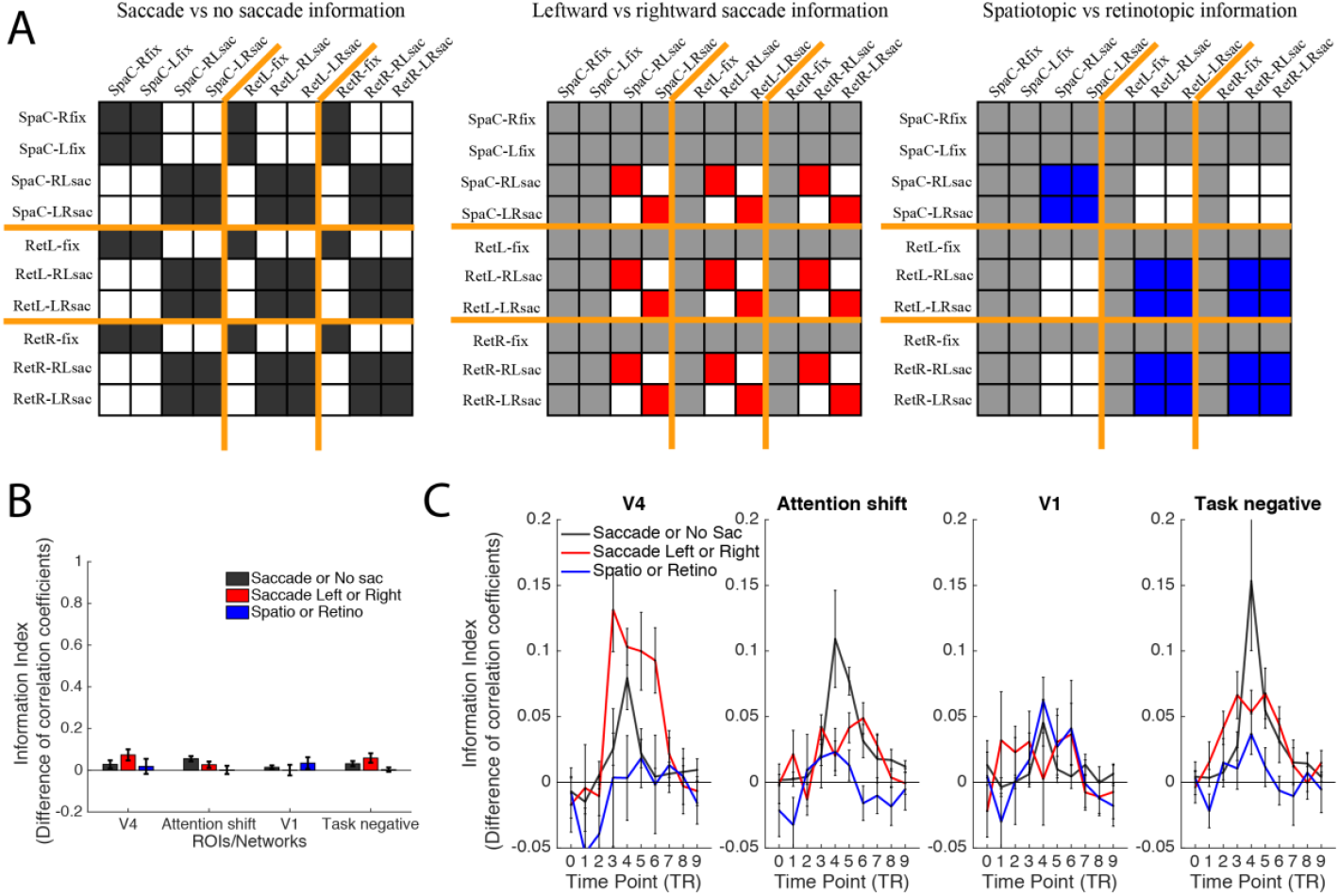
MVPA and MVPTC analyses and results of the Eyes-move task. (A) Hypothetical matrices for saccade vs no saccade, leftward vs rightward saccade, and spatiotopic vs retinotopic attention information. Orange lines separate conditions in spatiotopic (“attend center”), retinotopic left (“attend left of cross”), and retinotopic right (“attend right of cross”) blocks. Cells colored in dark grey, red, and blue are the within-group correlations, and white cells are the between-group correlations. Light grey cells are not used in the corresponding analysis. The index value of each type of information is calculated by subtracting the z-scored between-group correlation coefficients from the z-scored within-group correlation coefficients. (B) The index value of each type of information in each ROI/network; the scale is the same as Figure 5B. (C) The index value of each type of information at 10 time points, in each ROI/network; the scale is different from Figure 5B, 5C, and 6B. Error bars represent SEM. Extended Analyses are shown in Figure 6-1, 6-2, and 6-3 at the end.

**Figure 7.**
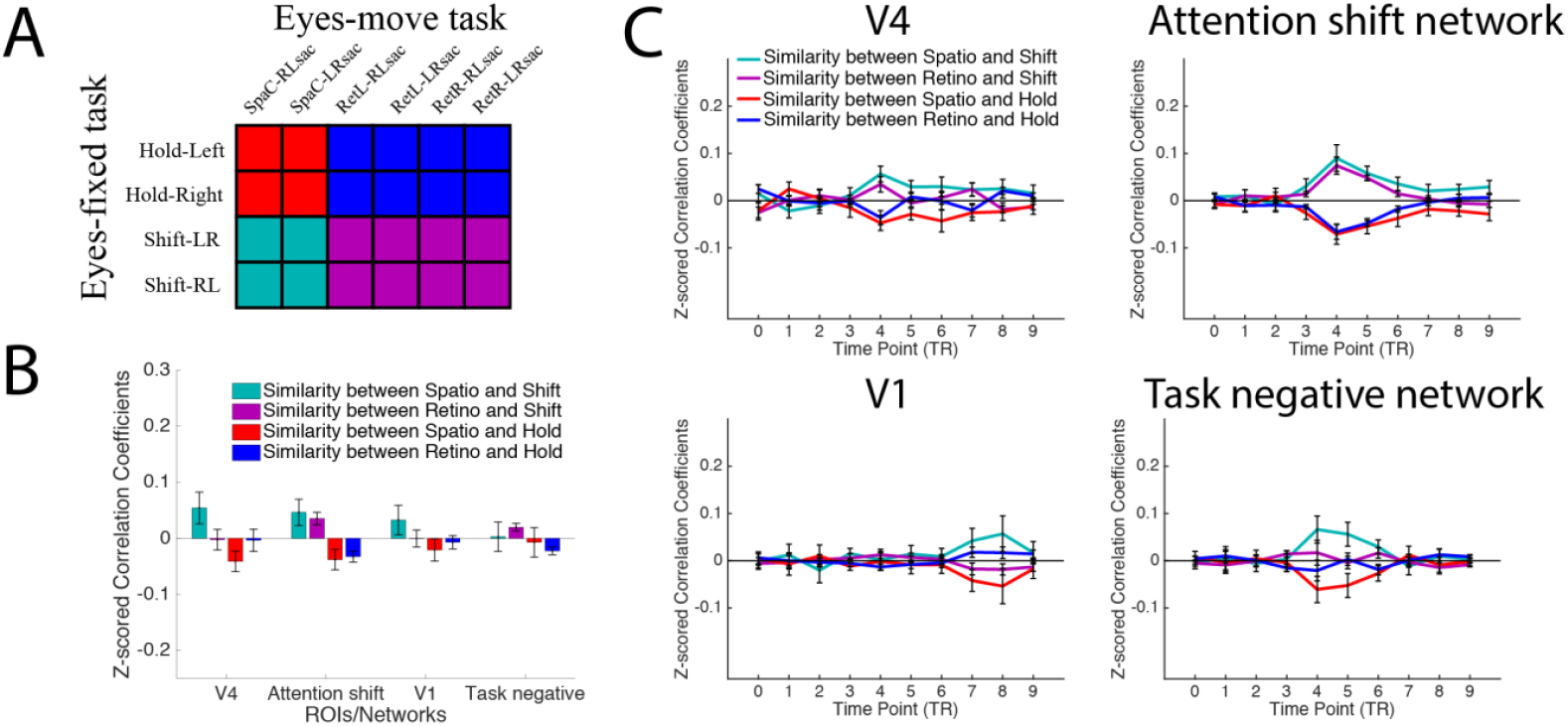
Cross-task similarity analyses in a priori ROIs/networks. (A) A hypothetical matrix indicating each combination of similarity: Retinotopic vs hold (blue), retinotopic vs shift (magenta), spatiotopic vs hold (red), and spatiotopic vs shift (cyan). (B-C) Pattern similarity (z-scored correlation coefficients) for each combination of conditions, for each ROI/network. (B) Pattern similarity based on whole-trial beta weights. (C) Pattern similarity timecourses based on FIR beta weights for each of 10 time points. Error bars represent SEM. Note that the roughly symmetrical patterns of the timecourse plots are likely due to the de-meaning step of subtracting the grand mean activity across conditions for each timepoint’s MVPA analysis (see Methods).

The results of this analysis are shown in Figure 7B & 7C and Table 3. In the whole-trial MVPA analysis, there was a significant main effect of similarity-to-shift versus similarity-to-hold in both V4 and the attention shift network, in that the representational similarity scores were generally higher when correlating both Eyes-move conditions with the shift-attention condition, compared to with the hold-attention condition. This main effect was significant at critical timepoint TP4 in both ROIs, and also at the neighboring timepoints TP3 and TP5 in the attention shift network. Interestingly, we did not find any significant main effect of Eyes-move condition for the whole-trial or for any timepoint that passed multiple corrections, nor were there any significant interactions in these two ROIs. The MVPTC plots indicate that both retinotopic and spatiotopic attention were more similar to shifting attention than to holding attention at the critical timepoint. In the attention shift network, the cross-task similarity patterns looked very similar for both reference frames, with both showing a strong effect of greater similarity to shifting attention. In V4, the similarity-to-shift effect seemed to be at least numerically greater for the spatiotopic attention condition, but this interaction was not significant. The main effect of similarity to Eyes-fixed was also significant (but of negligible magnitude) in V1 at TP3 and in the task negative network at TP4 and TP5.

**Table 3.** Statistics of 2×2 repeated-measure ANOVAs for each ROI at TP3, TP4, and TP5 respectively, on pattern similarity between Eyes-fixed conditions (hold & shift attention) and Eyes-move conditions (spatiotopic & retinotopic attention), separately for whole-trial analyses and time points of interest.

|  | Main effect of similarity to Eyes-fixed conditions (hold & shift) | Main effect of Eyes-move conditions (spatiotopic & retinotopic) | Interaction between similarity to Eyes-fixed conditions and similarity to Eyes-move conditions |
| --- | --- | --- | --- |
| V4 | $F=8.367, p=.015, \eta_p^2=.432^{**}$<br>TP3: $F=2.549, p=.139, \eta_p^2=.188$<br>TP4: $F=13.113, p=.004, \eta_p^2=.544^{**}$<br>TP5: $F=4.269, p=.063, \eta_p^2=.280$ | $F=0.486, p=.500, \eta_p^2=.042$<br>TP3: $F=0.510, p=.490, \eta_p^2=.044$<br>TP4: $F=0.922, p=.358, \eta_p^2=.077$<br>TP5: $F=0.184, p=.676, \eta_p^2=.016$ | $F=1.672, p=.223, \eta_p^2=.132$<br>TP3: $F=0.182, p=.678, \eta_p^2=.016$<br>TP4: $F=0.727, p=.412, \eta_p^2=.062$<br>TP5: $F=3.407, p=.092, \eta_p^2=.236$ |
| Attention shift network | $F=18.892, p=.001, \eta_p^2=.632^{**}$<br>TP3: $F=15.604, p=.002, \eta_p^2=.587^{**}$<br>TP4: $F=15.293, p=.002, \eta_p^2=.582^{**}$<br>TP5: $F=29.311, p<.001, \eta_p^2=.727^{**}$ | $F=0.291, p=.601, \eta_p^2=.026$<br>TP3: $F=0.291, p=.600, \eta_p^2=.026$<br>TP4: $F=5.514, p=.039, \eta_p^2=.334^*$<br>TP5: $F=0.827, p=.383, \eta_p^2=.070$ | $F=0.091, p=.768, \eta_p^2=.008$<br>TP3: $F=0.640, p=.441, \eta_p^2=.055$<br>TP4: $F=0.348, p=.567, \eta_p^2=.031$<br>TP5: $F=0.351, p=.566, \eta_p^2=.031$ |
| V1 | $F=3.606, p=.084, \eta_p^2=.247$<br>TP3: $F=12.968, p=.004, \eta_p^2=.541^{**}$<br>TP4: $F=0.575, p=.464, \eta_p^2=.050$<br>TP5: $F=1.469, p=.251, \eta_p^2=.118$ | $F=3.456, p=.090, \eta_p^2=.239$<br>TP3: $F=0.109, p=.748, \eta_p^2=.010$<br>TP4: $F=0.017, p=.898, \eta_p^2=.002$<br>TP5: $F=0.880, p=.368, \eta_p^2=.074$ | $F=0.473, p=.506, \eta_p^2=.041$<br>TP3: $F=0.232, p=.640, \eta_p^2=.021$<br>TP4: $F=0.229, p=.641, \eta_p^2=.020$<br>TP5: $F=0.026, p=.875, \eta_p^2=.002$ |
| Task negative network | $F=1.438, p=.256, \eta_p^2=.116$<br>TP3: $F=1.534, p=.241, \eta_p^2=.122$<br>TP4: $F=8.418, p=.014, \eta_p^2=.434^{**}$<br>TP5: $F=9.492, p=.010, \eta_p^2=.463^{**}$ | $F=0.397, p=.542, \eta_p^2=.035$<br>TP3: $F=0.012, p=.914, \eta_p^2=.001$<br>TP4: $F=4.807, p=.051, \eta_p^2=.304$<br>TP5: $F=3.332, p=.095, \eta_p^2=.232$ | $F=0.260, p=.620, \eta_p^2=.023$<br>TP3: $F=0.232, p=.639, \eta_p^2=.021$<br>TP4: $F=1.026, p=.333, \eta_p^2=.085$<br>TP5: $F=2.468, p=.144, \eta_p^2=.183$ |

\* indicate statistical significance at $p<.05$
\*\* indicate statistical significance at $p<.05$ (Holm-Bonferroni corrected for multiple post hoc comparisons, separately across ROIs/networks for whole-trial beta weights, and across three TPs for time-course beta weights)

While we did not find any significant interaction indicating a difference in representational similarity for retinotopic vs spatiotopic attention in our a priori ROIs/networks, are there other areas in the brain that might show differential similarity patterns? We next performed a searchlight analysis for a significant interaction effect at the critical time point TP4, as described in Methods. This analysis allowed us to extract potential regions showing one of two interaction patterns: (1) retinotopic relatively more similar to hold, spatiotopic relatively more similar to shift; or (2) spatiotopic relatively more similar to hold, retinotopic relatively more similar to shift. The searchlight revealed four clusters (Figure 8A and Table 4), all with the retinotopic-hold/spatiotopic-shift pattern. The clusters were located in ventral areas and superior parietal regions bilaterally, which were in later visual hierarchy in both ventral and dorsal pathways. No regions with the spatiotopic-hold/retinotopic-shift pattern survived the cluster threshold correction.

**Figure 8.**
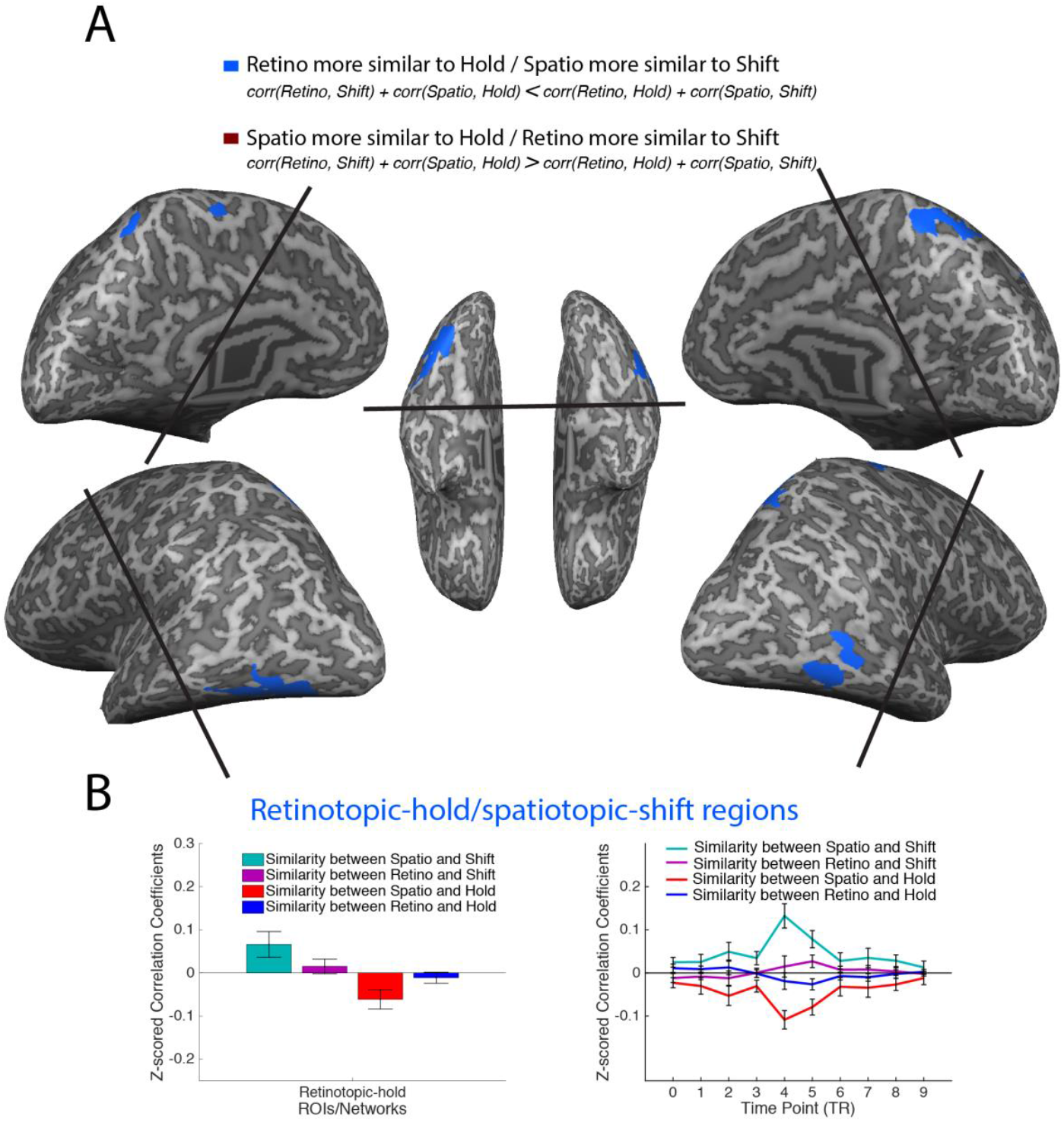
Cross-task pattern similarity, whole-brain searchlight analyses. (A) Regions showing a significant interaction effect in the searchlight. Regions exhibiting a significant retino-hold/spatio-shift pattern shown in blue; Regions exhibiting a significant spatio-hold/retino-shift pattern shown in scarlet (no clusters passed significance threshold for this contrast). Result is based on cross-task MVPTC, using the pattern correlation difference at TP4, with direction of contrast as indicated in the legend. The searchlight map was corrected for cluster-threshold in the same way as other brain maps. Searchlight analyses were conducted on the volume maps and projected onto an inflated brain for visualization purpose. (B) Pattern similarity timecourses (as in Figure 7) for each combination of conditions, shown for the retino-hold/spatio-shift areas extracted from A (all voxels averaged into single network; for separate plots for each individual area, see Figure S3). Plots are for illustrative purposes only to explore the specific pattern driving the significant interaction. Error bars represent SEM. Extended Analyses are shown in Figure 8-1 at the end.

**Table 4.** Description of clusters in regions with the retinotopic-hold pattern, including Talairach coordinates of the peak voxel, number of voxels, and t values.

| Networks | Areas | Hemisphere | TAL coordinates of peak voxel |  |  |  |  |
| --- | --- | --- | --- | --- | --- | --- | --- |
|  |  |  | x | y | z | # of voxels | t values (df=11) |
| Retinotopic-hold | Parahippocampal Gyrus | R | 39 | -49 | 4 | 691 | 4.0403 |
|  | Fusiform Gyrus | L | -37 | -59 | -12 | 708 | 4.9133 |
|  | Precuneus | R | 11 | -60 | 67 | 884 | 3.9127 |
|  | Paracentral Lobule | L | -3 | -41 | 60 | 472 | 3.4624 |

For illustration purposes, we plot the cross-task similarities for the regions identified in the searchlight (Figure 8B; plots for separate clusters in Extended data Figure 8-1). Note that this analysis is circular; we show the interaction patterns here for descriptive purposes only. The interaction in these regions seems to be primarily driven by the spatiotopic comparisons, particularly the high similarity between spatiotopic and shifting attention.

## Discussion

In summary, we were able to uncover several signatures of covert attention during fixation and across saccades from the multivoxel activation patterns in various brain regions. First, pattern similarity results from within the Eyes-fixed task support the validity of our design and analyses. In the visual and attention shift areas we could decode which location the participants were holding attention at, even dynamically in the time course, consistent with existing findings that attention modulates the activity in visual areas (Desimone & Duncan, 1995) and pattern activities in shift-related areas can be used to decode attention in the left vs right hemifield (Gmeindl et al., 2016). In the Eyes-move task we could similarly decode which hemifield was being covertly attended both before and after the saccade (Extended data Figure 6-1). We could also reliably decode from the Eyes-fixed task whether a covert attention shift was executed in the middle of the trials, specifically at the critical timepoint TP4 which corresponds to the transient shift, consistent with time-course decoding results about shift execution with SVM in_Chiu, Esterman, Gmeindl, & Yantis, 2012.

The more novel contributions of our study come the Eyes-move task and the patterns of representational similarity in our cross-task analysis. In the Eyes-move task, information about saccade execution emerged at TP4 in V4, and at all of TP3, TP4, and TP5 in the attention shift network as well as the task negative network. On the other hand, we found little information about whether attention was being held in retinotopic vs spatiotopic reference frames in the attention shift network or any of our other ROIs. Instead, with the cross-task pattern similarity analysis, we found that both spatiotopic and retinotopic attention were represented more similarly to shifting compared to holding attention, especially in the attention shift network. In addition, our searchlight analysis revealed brain regions where spatiotopic attention was represented more similarly to shifting and retinotopic relatively more similarly to holding attention (“retinotopic-hold / spatiotopic-shift” regions), but we did not find any brain regions displaying the opposite pattern. Below we discuss how our study contributes to the existing literature of covert vs overt attention shifts, and a few ways in which these novel findings of pattern similarity are informative to understanding the mechanisms of covert attention across saccades.

### Overlap between overt and covert representational patterns: Possible accounts

In the Eyes-move task, we found MVPA information in the attention shift network about saccade execution: i.e., the neural activity patterns in the attention shift network differentiated trials on which a saccade was executed from trials on which participants remained fixated. As mentioned in the introduction, overt and covert attention have been found to involve overlapping brain areas (Beauchamp et al., 2001; Corbetta et al., 1998; de Haan et al., 2008; Nobre et al., 2000; Perry & Zeki, 2000). Our study differed from these studies in that the paradigm used in these previous studies typically involved overt attention shift tasks where the focus of top-down attention was maintained at the fovea, both before and after saccades. In these cases, the saccade execution and the shift in attended location required by the task are intertwined, and the similarity in brain activity between overt and covert attention shifts could have been related to having the same “goal” between the eyes and attention. In daily life, overt and covert attention often have the same goal in space, but there are cases where the saccade and attentional targets could have different goals; for example, when we need to flexibly extract a lot of information within a short period of time (e.g., looking for cars while still paying attention to the traffic signals at a busy crossroads). In our design, rather than participants attending to and performing tasks about the saccade target, they were asked to maintain attention covertly on a peripheral cued location, while completing an eye movement between other locations. This allowed us to disentangle the saccade execution from the allocation of top-down task-directed attention. The fact that we still found representations in the same brain regions about saccade execution and covert shift execution might further contribute to the literature supporting overlapping neural mechanisms of covert attention shifts and saccades (Beauchamp et al., 2001; Corbetta et al., 1998; de Haan et al., 2008; Nobre et al., 2000; Perry & Zeki, 2000).

That said, there are several possible interpretations of this overlap between representations of saccades and covert attention shifts in our task. Here we discuss a few non-mutually exclusive explanations for why the saccade and no-saccade trials may have produced differentiable activation patterns in the attention shift network – and why in these areas, saccade trials of both references frames may have had greater representational similarity to the covert shift-attention trials in the cross-task similarity analysis.

One reason could be that covert shifting of attention is directly involved in making a saccade; i.e., the execution of the saccade required a pre-saccadic shift of attention towards the saccade target, and this initial covert shift was what was driving the representational similarity to the covert shift-attention trials. It has been widely shown that shifts of covert attention precede saccade execution (Godijn & Pratt, 2002; Peterson et al., 2004), and pre-saccadic attention is considered critical for determining the saccade endpoints to execute accurate saccades and enhancing perceptual representations of the saccade target (Gersch et al., 2004; Zhao et al., 2012). Even when the task is designed as attending to peripheral locations other than the saccade target, there is evidence that attention is still pre-saccadically shifted to the saccade target (Kowler et al., 1995). In our experiment, the information about saccade vs no saccade in the attention shift network emerged fairly early (around TP3), which could be related to the preparation stage (pre-saccadic shift stage) before the saccade was executed, potentially providing indirect support for this account.

Another potential account is that the task involved a covert shift of attention not related to execution of the saccade per se, but due to perisaccadic updating or remapping of the peripheral focus of attention, on both retinotopic and spatiotopic saccade trials. Previous studies involving spatiotopic remapping have found anticipatory remapping signals in the lateral intraparietal sulcus in monkeys (Duhamel et al., 1992), which could overlap with our parietal attention shift regions in humans. As described earlier, retinotopic attention can be seen as shifting attention relative to the screen/world, and spatiotopic attention can be seen as shifting attention relative to the eyes. One possibility is that both types of attention in our task involved some updating process across saccades that engaged an attentional shift signal in this brain region, which would be consistent with our cross-task correlation results that both spatiotopic and retinotopic attention were more similar to shifting compared to holding attention in the attention shift network. We discuss the implications of this aspect more in the section on retinotopic vs spatiotopic attention below.

A third possibility could be that our Eyes-move task may have triggered a more generic temporary disengaging/reengaging of top-down attention; i.e. a transient change or shift of attention on saccade trials that might have occurred independently of saccade planning, executing, or remapping processes. For example, although our task and instructions were designed to encourage continuous attention, we cannot rule out the possibility that participants may have approached the task as a serial attention task (attend the relevant stream, then disengage to execute saccade, then reengage again on the relevant stream), instead of attending continuously on the relevant stream. Or the abrupt onset of the saccade cue might have captured attention and caused an involuntary shift of attention away from the to-be-attended location. In cases like these examples, a transient shift in attention may have evoked representationally similar patterns of activity in this region on saccade trials to the goal-directed shifts of covert attention on fixation trials, without being directly related to the saccade itself. However, note that while we can’t rule out this third account, it is unlikely that this scenario could have accounted for our full pattern of results, particularly the searchlight findings.

While we cannot differentiate between the relative contributions of these mechanisms, our results nonetheless support a close link between the neural mechanisms associated with covert attention shifts during fixation and retinotopic/spatiotopic attention across saccades in the attention shift network. We also found additional similarities between representational patterns in area V4. In comparing the relative amounts and types of information present in the attention shift network versus area V4 patterns, we found an intriguing parallel; the attention shift network had relatively more information about the execution of covert attention shifts and saccades, while V4 had more information about the location of covert attention and the direction of saccades. This pattern aligns with the general understanding that the attention shift network is more involved in the *execution* of shifting spatial attention, and V4 in the *modulation* of spatial attention (Yantis et al., 2002).

Outside the domain of perisaccadic processing, previous literature has shown that the attention shift network is associated with broad, domain-independent brain activity for transient shifts of attention (Chica et al., 2013; Gmeindl et al., 2016; Greenberg et al., 2010; Shomstein & Yantis, 2004, 2006; Yantis et al., 2002). Our findings comparing covert attention shifts with attention updating across saccades further indicate that the brain activity patterns associated with covert attention shifts may be widely and reliably involved in various domains, contexts, and tasks.

### Representational patterns for retinotopic versus spatiotopic attention

How spatial attention is maintained/updated in particular reference frames across saccades has been an open question in the literature, and it is actively debated with various paradigms whether one reference frame is more native or dominant – and thus requires less updating across saccades – than the other (Crespi et al., 2011; Fabius et al., 2016; Fairhall et al., 2017; Golomb et al., 2008; Golomb & Kanwisher, 2012a, 2012b; Melcher & Morrone, 2003; Satel et al., 2012; Shafer-Skelton & Golomb, 2017; Turi & Burr, 2012; Zimmermann et al., 2013). In the case of spatial attention, it has been argued that attention pointers proactively remap to compensate for saccades and maintain spatiotopic attention (Cavanagh et al., 2010; Rolfs et al., 2011), but also that attention might linger in retinotopic coordinates even after a saccade (Golomb, 2019; Golomb et al., 2008, 2010; Jonikaitis et al., 2012; Marino & Mazer, 2018). More generally, spatiotopic remapping signals have been found in several brain regions, including monkeys’ lateral intraparietal area (LIP) (Duhamel et al., 1992), superior colliculus (SC) (Walker et al., 1995), frontal eye field (FEF) (Umeno & Goldberg, 1997), and striate and extrastriate cortex (Nakamura & Colby, 2002), and human visual and parietal cortex (Merriam et al., 2003, 2007). Higher-level visual and parietal areas in particular have also been a focus of much debate over dominant reference frames for neuronal receptive fields (Duhamel et al., 1997; Snyder et al., 1998), fMRI adaptation (Baltaretu et al., 2018; Fairhall et al., 2017; McKyton & Zohary, 2007; Zimmermann et al., 2016), functional organization (Crespi et al., 2011; d’Avossa et al., 2007; Golomb & Kanwisher, 2012b; Ward et al., 2010), and attentional modulation (Golomb et al., 2010; Rawley & Constantinidis, 2010).

In the current study, we approached this question from a different angle; to our knowledge, it is the first attempt to directly compare the brain activity patterns of covert attention maintained/updated in the periphery across saccades and during fixation. We found that in the pre-defined attention shift network, both retinotopic and spatiotopic attention evoked more similar representational patterns to covertly shifting attention than to covertly holding attention. Perhaps this is not surprising, given that both retinotopic and spatiotopic trials involved an eye movement, which is expected to engage attentional shifts as discussed above. In that sense, it is less notable that both retinotopic and spatiotopic resembled shifts more than holds per se; but the lack of a *relative* difference in representational similarity is intriguing. If attention were represented more natively in one reference frame, we may have predicted the other condition to show relatively more similarity to the shift condition. Our exploratory searchlight analysis did reveal some regions where spatiotopic attention was relatively more similar to shifting attention, but no regions with the opposite pattern. This asymmetry may reflect the idea that retinotopic attention is the more “native” coordinate system for spatial attention (Golomb et al., 2008), though it is interesting that neither this pattern nor the opposite pattern was found within the attention shift network itself.

In general, we found less of a difference between retinotopic and spatiotopic conditions than what might have been predicted. In analyses directly comparing the two reference frames, we did not reveal any representational difference between retinotopic and spatiotopic conditions in the whole-trial MVPA in the attention shift network or other ROIs, and in the MVPTC analyses, significant information about retinotopic vs spatiotopic attention was only found in V1 at TP4, but not in other pre-defined ROIs/networks or timepoints. In a series of extended analyses, we further probed for retinotopic vs spatiotopic differences. A whole-brain MVPTC searchlight revealed a few scattered regions outside our pre-defined ROIs/networks that showed information about spatiotopic vs retinotopic attention at individual time points (Extended data Figure 6-2). We also performed a whole-brain univariate contrast, and only found a small area in the right IPL where there was a univariate difference between spatial attention in retinotopic and spatiotopic reference frames across saccades (Extended data Figure 6-3). These results – along with the accurate behavioral task performance for both conditions – confirm that participants were allocating attention properly and differently in the two tasks; but given that reference frame conditions were blocked and participants knew which reference frame to attend to before each trial started, the interpretation of these direct spatiotopic vs retinotopic comparisons is more ambiguous.

Why didn’t we find greater differences in retinotopic vs spatiotopic patterns in our attention-related ROIs? One important consideration is that our task was designed to equate visual input across these two conditions. Both conditions contained constant, dynamic stimulation (RSVP streams) in the same three locations; the only difference was which of the streams, depending on which reference frame, was *attended* at any moment in time. This design is in contrast to a design commonly used in prior studies probing other aspects of reference frames across saccades, where only one stimulus is presented at a time, and retinotopic and spatiotopic conditions differ in terms of both stimulus-driven visual input and attentional locus (e.g. Baltaretu et al., 2018; Crespi et al., 2011; d’Avos sa et al,2007; fairhall et al., 2017; Gardner et al., 2008; Golomb & Kanwisher, 2012; McKyton & Zohary, 2007; Pertzov et al., 2011; Rawley & Constant midis, 2010; Zimmermann et al., 2016).

Moreover, our analysis was designed to look for representational signatures associated with attending in a retinotopic or spatiotopic reference frame (i.e., how shift- or hold-like they were); not to ask whether we could decode which particular retinotopic or spatiotopic locations were being attended. Early visual areas are known to be retinotopically organized (Crespi et al., 2011; d’Avossa et al., 2007; Gardner et al., 2008; Golomb & Kanwisher, 2012b; Merriam et al., 2013; Sereno et al., 1995), and we would expect that at least in these areas, attending to a particular retinotopic location across a saccade would look more similar to holding covert attention at that same retinotopic location during fixation than to shifting attention to a different retinotopic location (i.e., the brain activity pattern of RetL-RLsac would be more similar to Hold-L compared to Shift-LR, for example). Indeed, we could decode which hemifield(s) were attended on saccade trials (Extended data Figure 6-1), but this was not the goal of our study. Instead, the primary goal of the current study was to ask more broadly, whether the neural processes associated with maintaining attention in retinotopic (or spatiotopic) coordinates evoked more similar representational patterns to holding compared to shifting covert attention. Thus our analysis included correlations of conditions with both hemifields (e.g., similarity between retinotopic and hold attention includes correlations between RetL (with both RL and LR saccades) vs Hold-L, RetL vs Hold-R, RetR vs Hold-L, and RetR vs Hold-R; same for other cross-task correlations). This likely explains why we did not find a Retinotopic-Hold / Spatiotopic-Shift effect with the cross-task similarity searchlight analysis in early visual areas.

Instead, our cross-task pattern similarity analysis was better suited to reflect potential connections between the representations of covert attention across saccades and during fixations, independent of potential confounds from visual stimulation and hemifield-based attentional effects. Thus, it is telling that our pre-defined ROIs – particularly the attention shift network – did not show a difference in representational similarity between the retinotopic and spatiotopic reference frames in the cross-task similarity analysis, such that both were more representationally similar to shifting attention; but the exploratory searchlight analysis revealed some potential regions where spatiotopic attention was relatively more shift-like than retinotopic attention. As introduced earlier, eye movements distinguish the two reference frames in a way that retinotopy can be considered as “holding” a location relative to the eyes and “shifting” relative to the world, and spatiotopy can be considered as “shifting” relative to the eyes and “holding” relative to the world. One possibility is that maintaining spatiotopic and retinotopic attention across saccades may involve different types of updating that might be represented with “hold” and “shift” signals combined across different sets of regions. In other words, both types involve shift signals in the attentional shift network, whereas spatiotopic attention requires additional shift signals in other areas. That these other areas include bilateral anterior ventral areas and superior parietal regions, located in later visual hierarchy in both ventral and dorsal pathways, may hold further clues for understanding this complex process. In summary, coordination between different brain networks/regions may support more flexible updating of attention across saccades in different contexts, raising interesting follow-up questions regarding how and when this process might be achieved mechanistically, and how it is related to behavior, development, and clinical implications.

---

**Figure 6-1.**
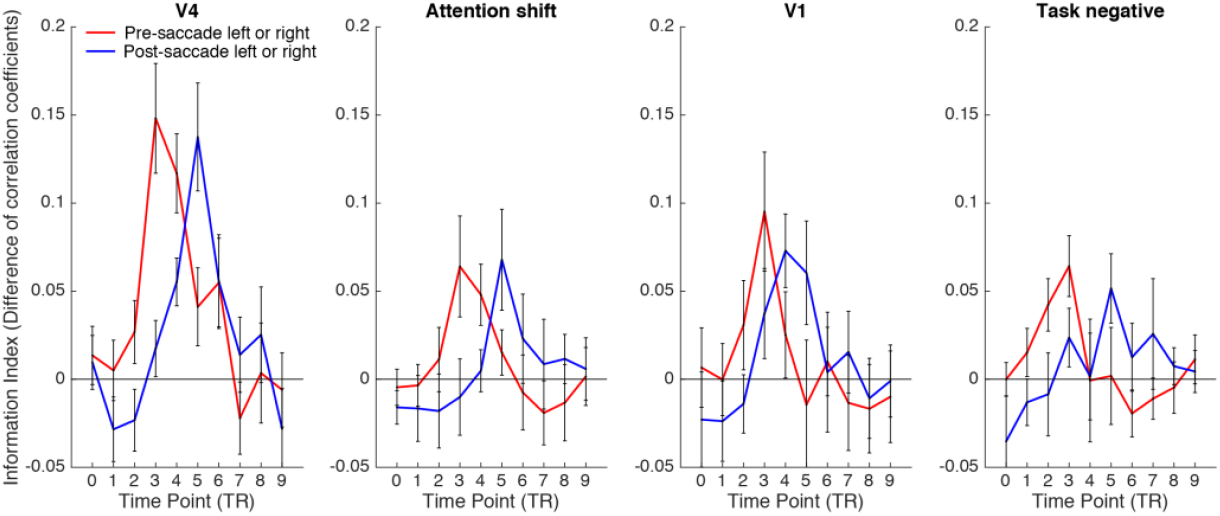
Extended Data showing MVPTC results of information about the hemifield attended (left or right) before and after the saccade separately (in the Eyes-move task). The index values of each type of information at 10 time points are plotted for each ROI/network. Error bars represent SEM. Results show that we could decode which hemifield was being covertly attended both before and after the saccade.

**Figure 6-2.**
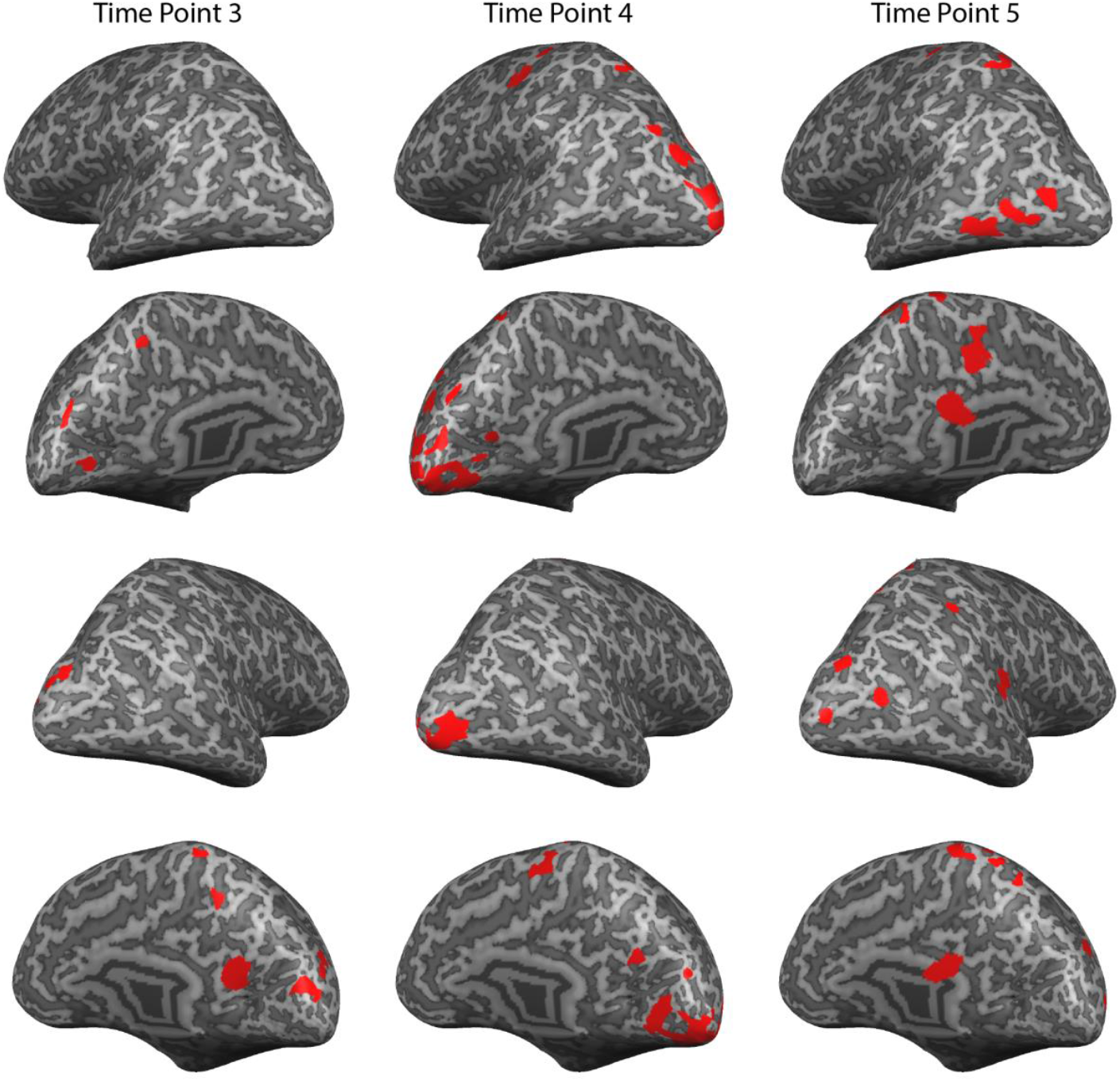
Extended Data showing information of retinotopic vs spatiotopic attention in searchlight analyses, for time point 3, 4, and 5 separately. This whole-brain analysis is analogous to Figure 6, blue condition (information about spatiotopic vs retinotopic). Red areas show significant information after cluster-threshold correction at p<.05. The viewing angle for each row is left lateral, left medial, right lateral, and right medial, respectively.

**Figure 6-3.**
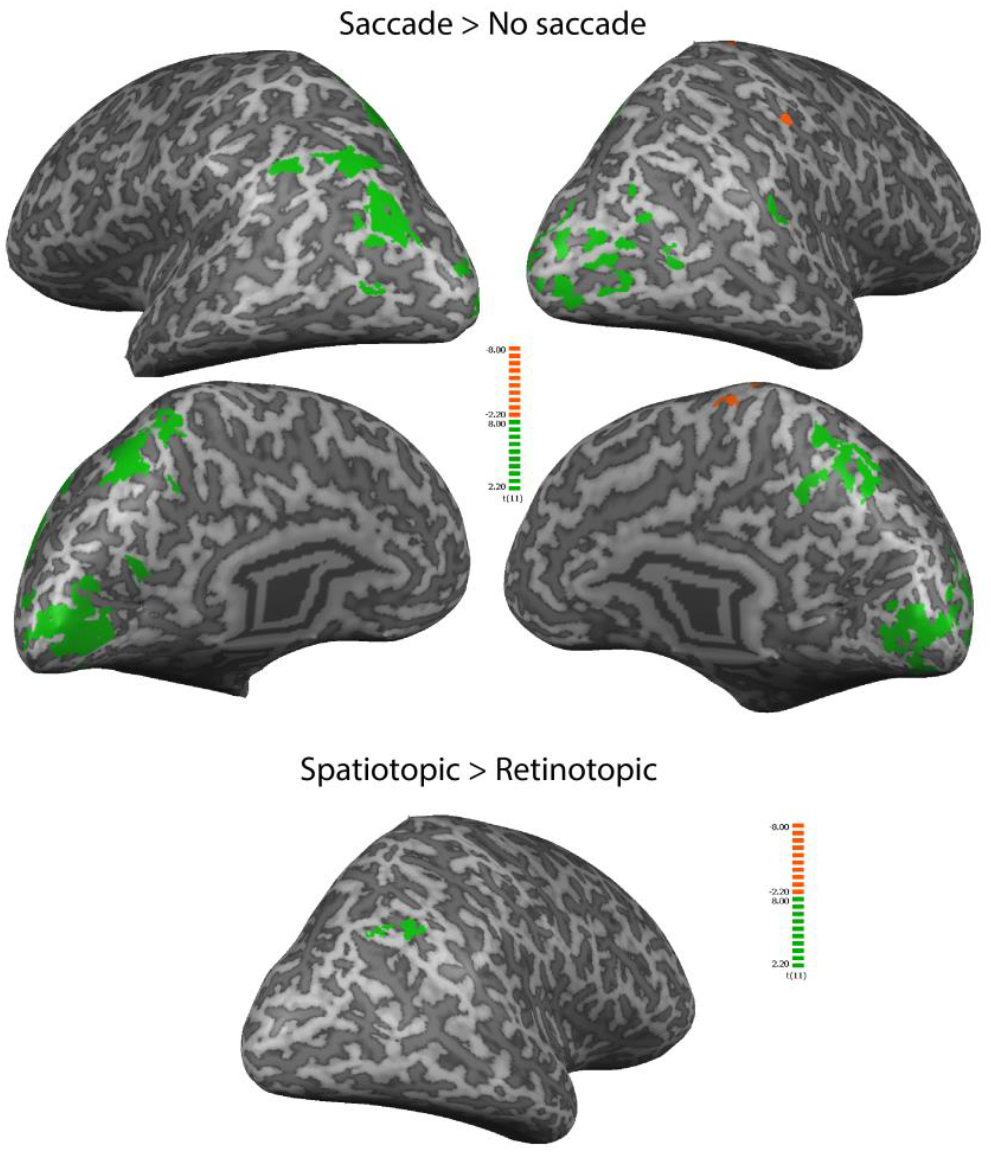
Extended Data showing univariate differences, based on whole-trial betas, between saccade and no saccade conditions in the Eyes-move task (top), and between retinotopic and spatiotopic conditions in the Eyes-move task (bottom). For each contrast, significant clusters in the positive direction are shown in green and negative in orange. Maps were cluster threshold corrected at p<.05. For Spatiotopic > Retinotopic contrast, the only significant cluster found was located in the left hemisphere, so only the left lateral viewing angle is shown here.

**Figure 8-1.**
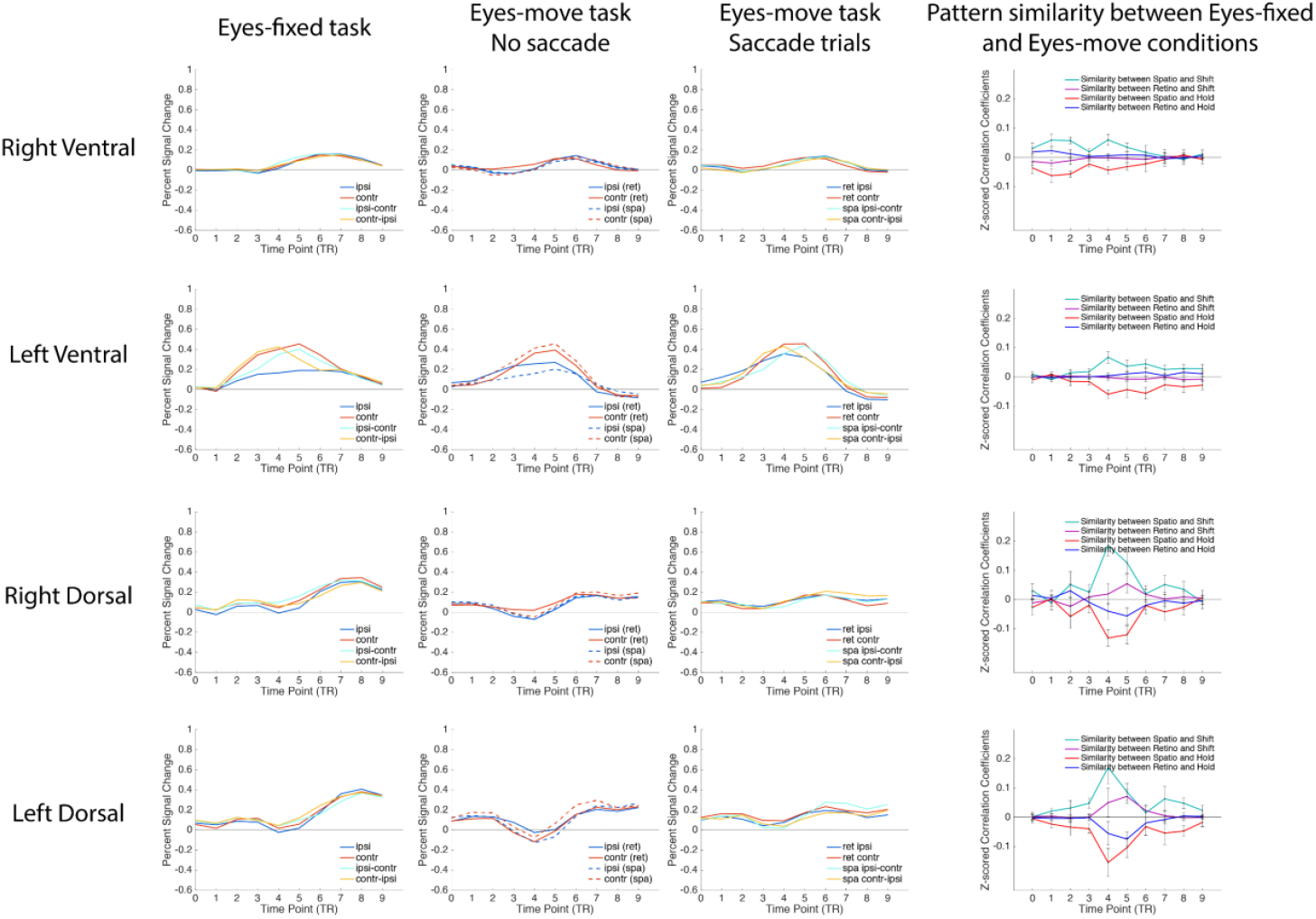
Extended Data showing univariate activation (the first three columns) and cross-task pattern similarities (the last column), separately for each cluster of the retinotopic-hold regions from the exploratory searchlight analyses. The univariate activation plots were comparable to Figure 4, and the pattern similarity plots to Figure 7C & 8B.

## Footnotes

1 Note that the attention shift network was defined by the univariate contrast of shift > hold (with the whole-trial betas), so these MVPA results are not completely independent, although a univariate effect alone (linear transform) could not drive a correlation-based MVPA difference. Nonetheless, the MVPTC results are useful as a validity check, and the remaining analyses that we focus on below are fully independent of the ROI definitions.

